# Multiscale Temporal Processing Supports Sound Recognition under Causal Constraints

**DOI:** 10.64898/2026.07.27.740874

**Authors:** Michele Esposito, Tonio Weidler, Christian Ferreyra, Bruno L. Giordano, Elia Formisano

**Affiliations:** Institut des Neurosciences de La Timone, CNRS UMR 7289 – Université Aix-Marseille, Marseille, France; Department of Cognitive Neuroscience, Faculty of Psychology and Neuroscience, Maastricht University, Maastricht, The Netherlands; Laboratoire d’Informatique et des Systèmes, UMR 7020, CNRS and Université Aix-Marseille, Marseille, France; Maastricht Centre for Systems Biology and Bioinformatics, Faculty of Science and Engineering, Maastricht University, Maastricht, The Netherlands

**Keywords:** Multiscale temporal processing, Causal sound recognition, Recurrent neural networks, Time-resolved representations, Computational auditory neuroscience

## Abstract

The auditory system operates under a fundamental computational constraint: at any moment, it has access only to past and present acoustic information. At the same time, it processes sounds across multiple temporal scales, although the computational advantage of this organization remains unclear. Despite both being defining characteristics of biological auditory processing, causal processing and multiscale temporal processing are rarely considered together in computational models. Here, we introduce the Multiscale Convolutional Recurrent Neural Network (MSCRNN) architecture, a brain-inspired model designed to test whether processing sounds across multiple temporal contexts improves recognition under causal constraints. Multiscale processing substantially improved recognition under causal constraints, enabling performance comparable to non-causal architectures after an initial evidence-accumulation period while providing only limited benefit when future acoustic information was available. Analyses of the network’s internal representations revealed a progressive transformation from stream-specific acoustic representations to increasingly integrated category-level representations across recurrent processing stages. Together, these findings provide a computational rationale for why biological auditory systems may benefit from multiscale temporal processing under causal constraints and establish the MSCRNN as an interpretable framework for generating and testing hypotheses about the temporal dynamics of auditory processing using electrophysiological and neuroimaging data.

## 1 Introduction

The auditory system continuously interprets sounds as they unfold in time, identifying acoustic events with widely varying temporal dynamics, from brief impacts to sustained environmental sounds such as rain, wind, or traffic. A fundamental challenge is that this process must operate causally: at any given moment, only past and present acoustic information is available. Consequently, recognition must rely on incomplete and continuously accumulating evidence, progressively refining perceptual interpretations as additional information becomes available. Despite their impressive performance, many contemporary deep learning models for environmental sound recognition process complete spectrograms or fixed-duration audio segments and generate predictions only after observing the entire acoustic signal (Hershey et al., 2017; Kong et al., 2020; Cakır et al., 2017; Esposito et al., 2024, e.g.). Neurocomputational models of auditory cortex have largely adopted the same assumption (Kell et al., 2018; Esposito et al., 2024, e.g.), limiting their ability to explain how sound representations evolve under causal constraints or how recognition emerges from the gradual accumulation of acoustic evidence. Beyond computational neuroscience, causal processing is also essential for real-time computer audition systems, where predictions must be generated while sounds are still unfolding.

Previous research has shown that the auditory cortex processes sounds across multiple temporal scales (Chi et al., 2005; Elhilali and Shamma, 2008; Poeppel, 2003; Boemio et al., 2005; Giraud and Poeppel, 2012; Ding and Simon, 2009; Woolley et al., 2005; Singh and Theunissen, 2004; Theunissen et al., 2000; Rupp et al., 2025). Shorter temporal scales support the analysis of transient acoustic events and fine temporal structure, whereas longer scales capture more slowly evolving acoustic regularities such as speech prosody and melodic contours (Santoro et al., 2014; Norman-Haignere et al., 2022). Neuroimaging studies further suggest that sensitivity to these temporal scales is organized across auditory cortex, with posterior-lateral regions responding preferentially to rapidly varying acoustic features and anterior-medial regions exhibiting greater sensitivity to slower temporal modulations (Santoro et al., 2014, 2017). This organization is further shaped by hemispheric asymmetries in spectro-temporal processing (Albouy et al., 2020). Electrophysiological recordings (Jasmin et al., 2019) and computational modelling studies (Zulfiqar et al., 2020, 2021) suggest that these large-scale functional differences may arise from systematic variations in neuronal temporal integration properties across cortical fields, potentially supporting the formation of abstract representations along the auditory processing hierarchy (Rauschecker and Scott, 2009).

Although multiscale temporal processing is well established in biological audition, its computational role under the causal constraints of real-time perception remains unclear. While causal neural architectures have been developed for tasks such as speech separation and enhancement (Luo and Mesgarani, 2018, 2019), they were not designed to investigate the role of multiscale temporal processing in environmental sound recognition. Because causal perception relies on continuously accumulating acoustic evidence, processing sounds simultaneously over multiple temporal contexts may provide a computational advantage over relying on a single temporal context. We therefore hypothesise that parallel processing across multiple temporal contexts improves sound recognition under causal constraints by reducing the ambiguity associated with incomplete acoustic evidence.

Testing this hypothesis requires an architecture that processes multiple temporal contexts while operating under strict causal constraints. We therefore introduce the *MultiScale Convolutional Recurrent Neural Network (MSCRNN)* architecture, a neuroconnectionist model (Doerig et al., 2023) comprising three parallel recurrent streams receiving different input temporal contexts. The architecture is inspired by the hierarchical temporal organization of the auditory cortex (Rupp et al., 2025; Norman-Haignere et al., 2022). By systematically contrasting causal and non-causal variants of the MSCRNN architecture with single-scale baselines, we show that processing acoustic information across multiple temporal contexts substantially improves sound recognition under causal constraints, whereas its benefit is markedly reduced when future acoustic information is available.

Beyond this performance-based comparison, we investigate how information is represented across the proposed architecture. Specifically, we ask whether individual temporal streams develop different representational profiles and whether combining information across streams yields representations that are more informative than those of any single stream alone. To address this question, we employ two complementary analyses. First, we use representational similarity analysis (RSA) (Kriegeskorte et al., 2008) to characterise how category-related representational geometry evolves across layers, streams, and time. Second, we examine how information about the temporal modulation information, a neurophysiologically grounded description of sound dynamics (Theunissen et al., 2000; Chi et al., 2005; Santoro et al., 2014, 2017), is represented and transformed throughout the network.

We make three main contributions. First, we show that multiscale temporal processing substantially alleviates the performance costs imposed by causal sound recognition, reducing the gap with non-causal architectures. Second, we show that multiscale processing supports a hierarchical transformation of sound representations, in which category-level organisation emerges alongside a reorganisation of temporal modulation information across the network hierarchy. Third, we introduce a biologically inspired multiscale recurrent architecture for causal sound recognition that is directly compatible with time-resolved electrophysiological analyses.

## 2 Results

To investigate the interaction between multiscale processing and causal sound recognition, we developed a MultiScale Convolutional Recurrent Neural Network (MSCRNN) architecture, a time-resolved architecture explicitly designed for online auditory processing (Figure 1). Unlike conventional sound recognition systems that process complete spectrograms offline, MSCRNNs generate framewise predictions every 10ms using only current and past acoustic information.

**Fig. 1:**
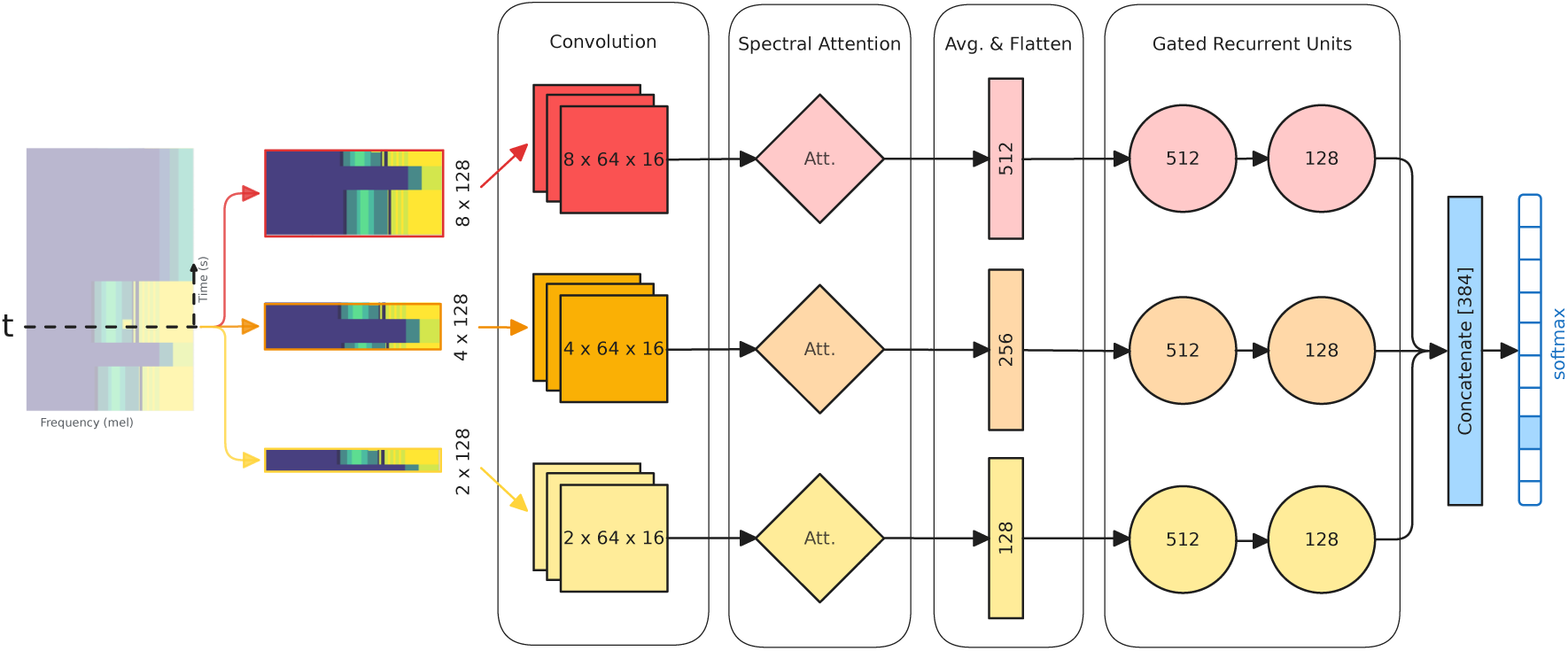
Model Architecture. The MSCRNN architecture comprises three parallel processing streams receiving different causal temporal contexts from a log-Mel spectrogram sampled at 10ms resolution. At time *t*, for example, the 40 ms stream the spectrogram segment spanning [*t −* 3*, t*]. Within each stream, convolutional, attention, and recurrent layers extract spectro-temporal features. The resulting stream-wise representations are concatenated to produce frame-wise sound category predictions.

The architecture combines multiple parallel streams operating over distinct input temporal contexts. Convolutional layers extract local spectro-temporal features within each stream, while recurrent dynamics progressively integrate information over time before combining stream-wise representations into a shared decision layer. Parallel temporal streams may allow the system to preserve both short-term acoustic detail and longer contextual structure without requiring access to future information. Detailed architectural specifications are provided in the Methods and Supplementary Methods Sections S-M2–S-M5 and Supplementary Tables S1–S7.

### 2.1 Multistream architectures outperform single stream networks

We first asked whether processing sounds across multiple temporal contexts improves recognition under causal constraints. To this end, we compared the multistream architecture with temporal contexts of 20–40–80 ms to all of its individual variants, as well as a 200ms single stream architecture that matches the multistream architecture in its number of parameters. Each architecture was trained independently 10 times. The 20–40–80 ms regime was determined by sequential hyper-parameter optimisation (Supplementary Methods Section S-M4; Supplementary Figure S6 and Supplementary Table S6).

Figure 2a show the framewise F1 scores, computed independently at each 10 ms time step by comparing the predicted and ground-truth sound labels for all of these model variants. The multistream architecture significantly outperformed all single stream architectures (non-overlapping bootstrap confidence intervals at *α* = 0.05). Additionally, amongst single stream models, larger context sizes (and therefore also larger numbers of parameters) do not imply better performance.

**Fig. 2:**
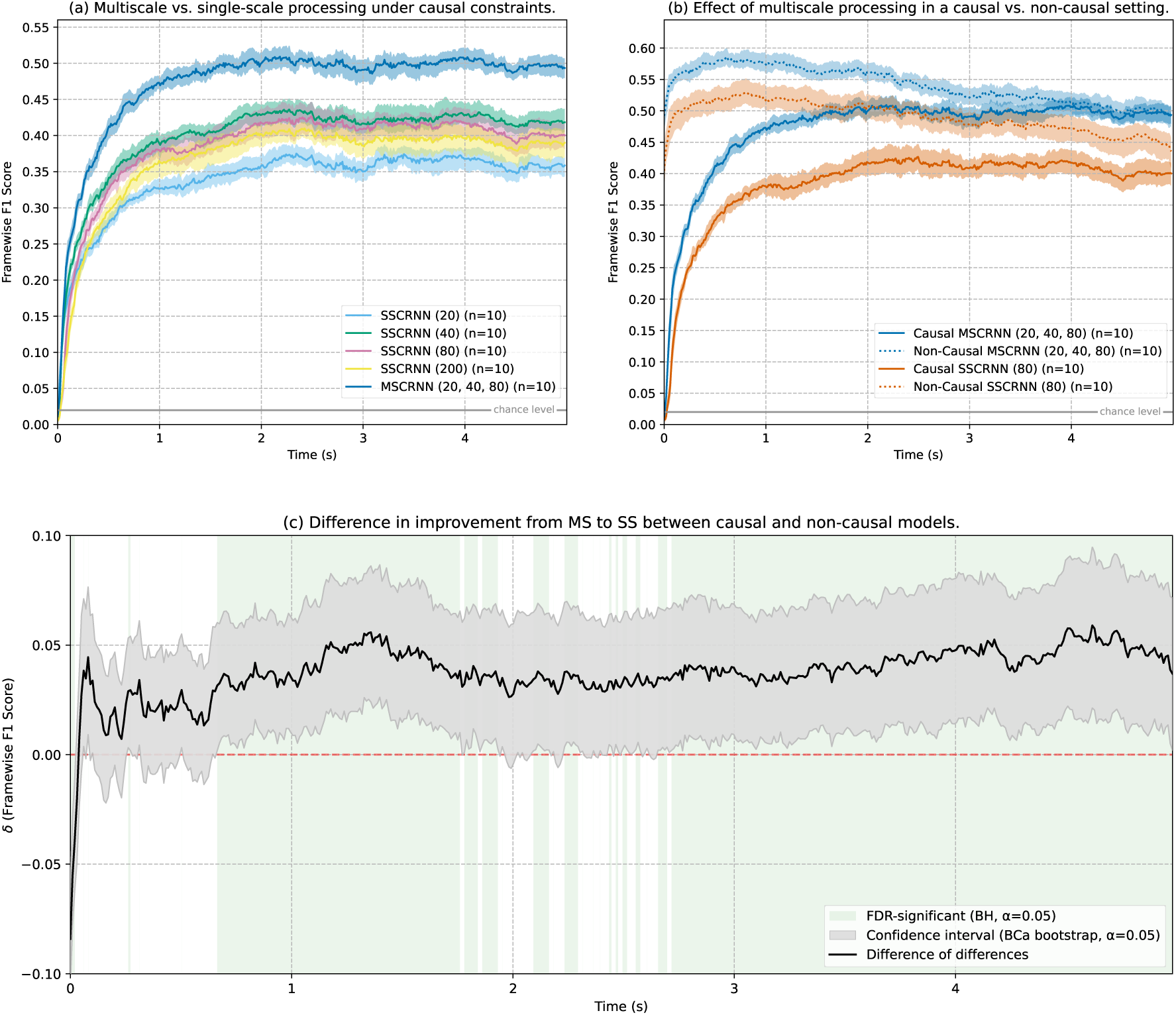
Framewise F1-scores of multistream (MS) and singlestream (SS) architectures, benchmarked under causal and non-causal settings. (a) Comparison of the 20–40–80 ms multistream regime to all its individual single stream variants under causal constraints and to a single stream model (SSCRNN-200) matched for number of parameters. The multistream configuration consistently outperforms the single stream components. (b) Comparison of a single-stream and a multistream model under causal (solid lines) and non-causal (dotted lines) conditions. Processing at multiple temporal scales has a substantially positive effect on performance under causal constraints, but not in the non-causal setting. (c) Interaction effect illustrating how much more multiscale processing benefits the model under causal constraints compared to the non-causal setting. Specifically, for four groups of models as shown in panel (b) of this figure, causal MSCRNN (cMS), non-causal MSCRNN (MS), causal SSCRNN (cSS), and non-causal SSCRNN (SS), we show *δ* = (*µ_cMS_ − µ_cSS_*) *−* (*µ_MS_ − µ_SS_*). Shaded green areas mark significant time points at FDR 0.05 (Benjamini-Hochberg procedure (Benjamini and Hochberg, 1995)) using bootstrap p-values from inverting confidence intervals. All results (a-c) report the means of 10 trainings on independent seeds per setting and shaded areas indicate the 95% bias-corrected and accelerated (Efron, 1987) bootstrap confidence intervals (*B* = 10, 000).

We additionally probed a regime with larger temporal context sizes (50–200–400) drawn from empirical neuroscience (Norman-Haignere et al., 2022). Results are reported in the supplementary material. The overall relationship between the multiscale network and its single-scale constituents remained the same. Importantly, larger temporal windows did not improve performance in neither the multiscale nor the single-scale architectures. In fact, in the multiscale condition, the design with shorter temporal scales reached higher F1 scores consistently over the full signal despite relying on substantially narrower input and convolutional kernels.

### 2.2 Multiscale processing is particularly useful under causal constraints

Next, we asked whether multiscale processing helps biological systems to alleviate the informational limitations posed by causality. To this end, we measured model performance (Figure 2b) of multiscale and single stream architectures with and without future context to explore the interaction effect.

Non-causal multistream models achieved high framewise F1 scores almost immediately after stimulus onset, reflecting their access to future acoustic information through bidirectional processing. In contrast, causal multistream models exhibited an initial evidence-accumulation period of approximately 2s before progressively approaching the performance of their non-causal counterparts. In both the causal and the non-causal setting, multistream models consistently outperformed the corresponding single-stream architecture. However, the performance advantage of multistream processing was markedly greater under causal constraints than under non-causal conditions. We confirmed this interaction by a difference-of-differences analysis (Figure 2c), which shows that the multistream advantage is significantly larger for causal than for non-causal architectures over a substantial portion of the sound.

Together, these findings indicate that multiscale processing provides a selective benefit under causal constraints, reducing the performance cost associated with the absence of future acoustic information.

### 2.3 Categorical separation emerges through hierarchical temporal integration

While the previous analyses established that multiscale processing improves recognition performance, it does not reveal how the individual temporal streams contribute to the model output. We therefore examined how categorical separation develops over time in the causal multiscale architecture (20–40–80 ms), comparing the full three-stream model (MSCRNN) with the corresponding single-stream models (SSCRNN; Figure 3).

**Fig. 3:**
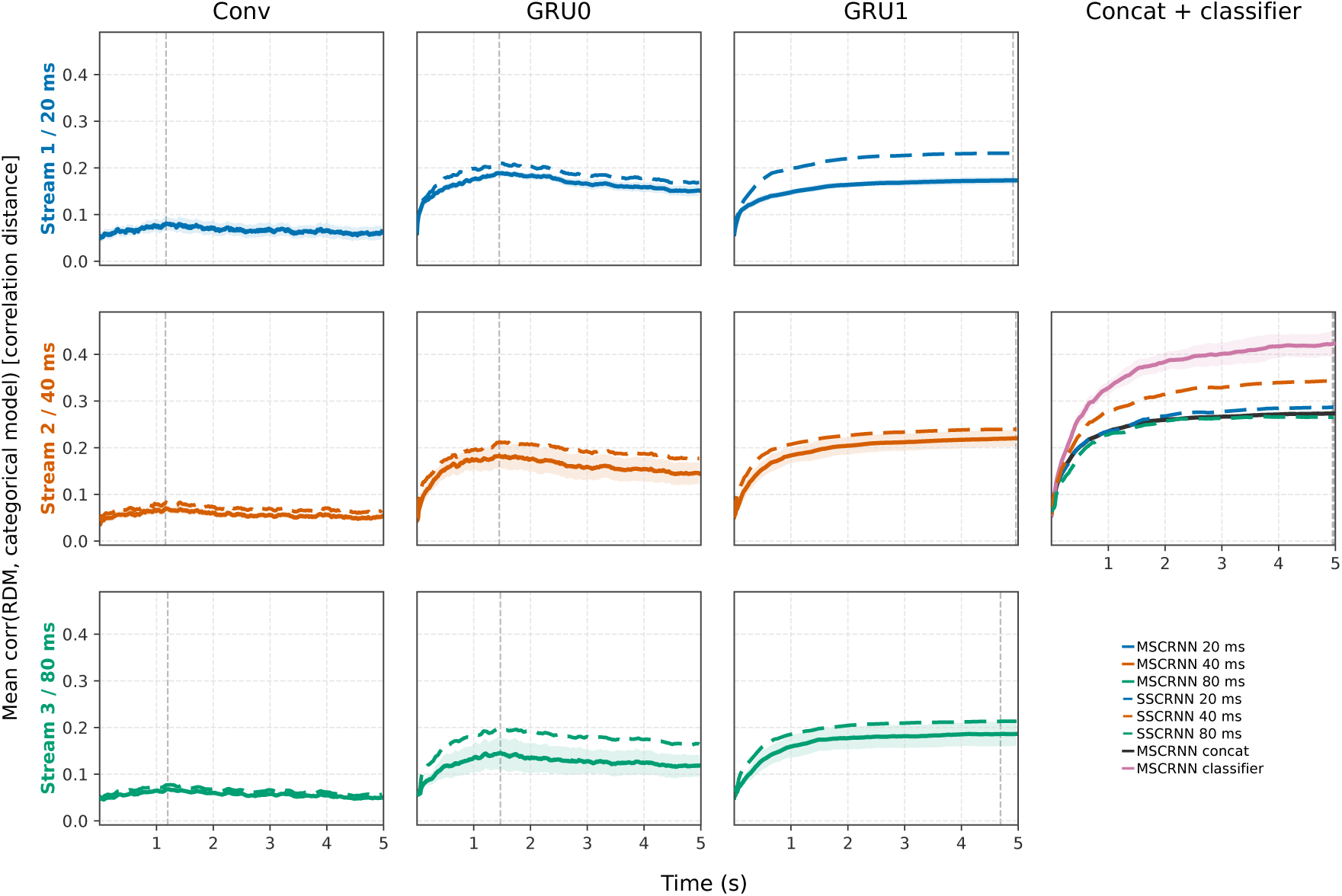
Time-resolved Pearson correlation between layer-wise representational dissimilarity matrices and the categorical model, averaged across three training runs. Rows: temporal streams (20, 40, 80 ms). Columns: Conv, GRU 0, GRU 1, and a combined late panel (concatenation, multistream classifier, single-stream classifiers). Solid lines: MSCRNN; dashed lines: SSCRNN. Shaded bands: *±*1 s.d. across runs (shown for both MSCRNN and SSCRNN). Vertical dashed lines: peak correlation time. Notice that: (i) Conv layers show weak correlation (mean *r ≈* 0.06, range 0.056–0.069). (ii) GRU layers increase correlation (mean *r ≈* 0.15–0.20 on MSCRNN streams); in per-stream GRU panels, the matching single-stream model wins in 6/6 comparisons (mean and peak). (iii) Stronger correlation appears after stream integration: MSCRNN concatenation mean *r* = 0.25 (peak = 0.27 at *∼*5 s); MSCRNN classifier mean *r* = 0.37 with peak *r* = 0.42 at *∼*5 s, above single-stream classifiers (peak *r* = 0.29, 0.34, and 0.27 for 20, 40, and 80 ms). (iv) The multistream advantage is mainly a fusion-plus-classification effect, not a uniform gain within every stream alone.

Across all streams, the convolutional layers showed only weak correlation with the categorical RDM (MSCRNN peak *r ≈* 0.07–0.08 at *∼*1.2 s), indicating that local spectro-temporal feature extraction alone provides little category-level separation. A marked increase emerged in the first recurrent stage (GRU0), where correlations rose rapidly and peaked early (*∼*1.4–1.5 s; MSCRNN Stream 20: *r* = 0.19; Stream 40: *r* = 0.18; Stream 80: *r* = 0.15). Individual SSCRNN reached comparable or slightly higher correlations at this stage (e.g. Stream 20: *r* = 0.21), suggesting that information on categorical separation can arise from temporal integration within a single temporal window. Categorical separation continued to develop in the second recurrent stage (GRU1), where correlations grew more gradually and peaked much later (*∼*4.7–5.0 s; MSCRNN Stream 40: *r* = 0.22; Stream 20: *r* = 0.17; Stream 80: *r* = 0.19). Again, individual SSCRNNs achieved correlations comparable or slightly exceeding those of the corresponding MSCRNN streams (e.g. Stream 40: *r* = 0.24), indicating that the benefit of multiscale processing is not expressed within isolated strams. Instead, the advantage of the multistream architecture became apparent after stream fusion. The concatenated representation reached a peak of *r* = 0.27 at *∼*5 s, while the final classifier achieved the strongest alignment (mean *r* = 0.37; peak *r* = 0.42 at *∼*5 s), exceeding all single-stream classifiers (peaks *r* = 0.29, 0.34, and 0.27 for 20, 40, and 80 ms models, respectively).

Together, these findings support a hierarchical progression of category-related representations. Category separation is weak in the convolutional layers, and becomes most pronounced after combining information across temporal streams. Notably, although individually trained single-stream models often achieve stronger category separation within their own representations than the corresponding MSCRNN streams, their final classifiers remain consistently inferior to the multistream model. This suggests that multistream training encourages individual streams to develop complementary representations whose value is realised primarily after combination rather than in isolation.

### 2.4 Representations across streams are predictive of differential temporal modulations

To investigate how acoustic information is distributed and integrated across temporal streams, we examined the extent to which the temporal modulation structure of sounds could be reconstructed from the hidden representations of individual streams and their combined activity. Reconstruction was performed separately for the first (GRU0) and second (GRU1) recurrent layers, the first stages of the network that explicitly integrate information over time, enabling us to characterize how temporal modulation representations evolve across the processing hierarchy.

Figure 4 shows reconstruction accuracy as function of temporal modulation rate and time for the three temporal streams (20 ms, 40 ms, and 80 ms) and for their combined representation. In GRU0, all three streams contained substantial information about the temporal modulation structure of the sounds, with reconstruction spanning a broad range of modulation rates throughout the sound. The 80 ms stream consistently yielded the strongest reconstruction, whereas the 20 ms and 40 ms streams showed slightly weaker but qualitatively similar patterns. The combined representation closely resembled the individual streams while providing a modest improvement in reconstruction accuracy, indicating that the first recurrent layer largely preserves acoustic information captured within each temporal context.

**Fig. 4:**
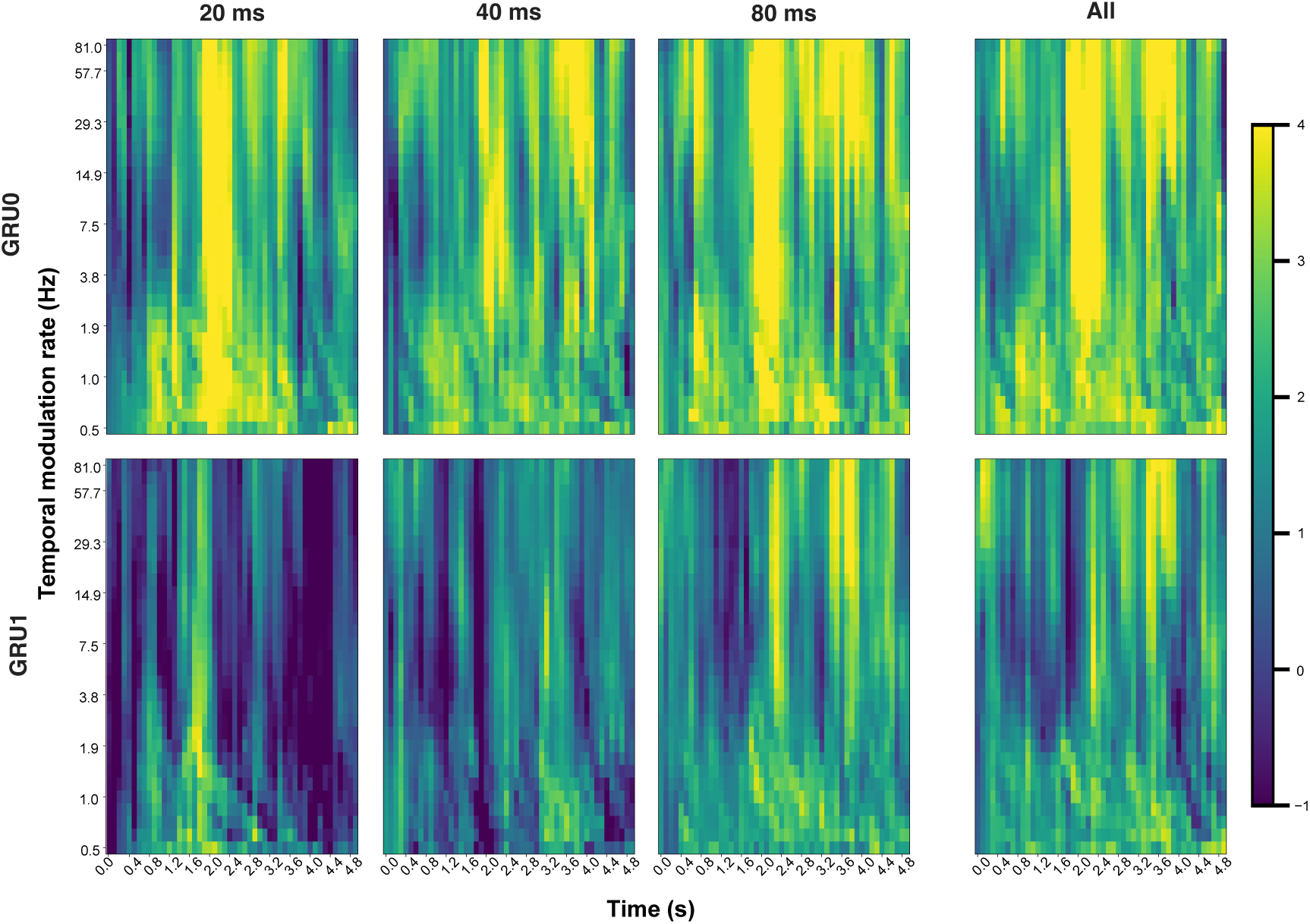
Reconstruction of temporal modulation features from recurrent representations. Reconstruction accuracy (z-scored relative to a permutation-based null distribution) for temporal modulation features across modulation rates and time. Columns correspond to individual temporal streams (20 ms, 40 ms, 80 ms) and their combined representation (All), while rows correspond to the first (GRU0) and second (GRU1) recurrent layers. In GRU0, substantial modulation information is present in all streams, with the 80 ms stream showing the strongest reconstruction and the combined representation closely resembling the longest stream. In GRU1, reconstruction from individual streams decreases, whereas the combined representation retains broader and more stable modulation information than any single stream. This pattern indicates a transition from stream-specific acoustic representations in GRU0 to integrated multiscale representations in GRU1.

A markedly different pattern emerged in GRU1. Reconstruction accuracy decreased substantially in the individual streams, particularly for intermediate and faster temporal modulation rates, indicating a progressive transformation away from detailed acoustic representations. In contrast, the combined representation retained considerably more temporal modulation information than any individual stream, with the improvement being most pronounced for slower modulation rates. Thus, although individual streams progressively lose acoustic detail, their combination preserves complementary information that remains accessible after integration.

Together, these findings reveal a hierarchical reorganization of temporal information within the network. The first recurrent layer largely preserves the temporal modulation structure present within each stream, whereas the second recurrent layer progressively transforms these representations while retaining complementary low-frequency modulation information after stream fusion. This transition coincides with the emergence of stronger category-level representations observed in the RSA analyses, suggesting that multiscale processing simultaneously supports increasing category selectivity and the preservation of behaviourally relevant slow temporal dynamics.

## 3 Discussion

In the present study, we investigated whether and how multiscale temporal analysis supports *causal* auditory recognition of natural sounds. To this end, we introduced a multiscale convolutional recurrent neural network (MSCRNN) architecture that performs framewise sound recognition at a temporal resolution of 10 ms while accessing only current and past acoustic information. Unlike most deep learning approaches to sound classification, which process complete spectrograms offline and generate a single prediction per sound (Esposito et al., 2024; Hershey et al., 2017; Cakır et al., 2017), MSCRNNs produce temporally evolving predictions as the acoustic signal unfolds.

The central finding is that multiscale processing aids causal sound recognition and that this benefit is selective compared to non-causal settings. Causal multistream models substantially outperform their single-stream counterparts and progressively approach the performance of non-causal architectures after an initial accumulation period. By contrast, introducing parallel temporal streams provides only a limited additional benefit when future acoustic context is already available. This interaction indicates that multiscale temporal processing is not merely a generic increase in architectural capacity, but a computational strategy that is particularly useful when recognition must operate online.

### 3.0.1 A Computational Account of Multiscale Processing under Causal Constraints

Our findings provide a computational rationale for multiscale temporal processing in biological audition. Previous work has established that auditory cortex analyses sounds across multiple temporal contexts, with different cortical regions exhibiting preferential sensitivity to acoustic information unfolding over distinct timescales (Chi et al., 2005; Giraud and Poeppel, 2012; Norman-Haignere et al., 2022; Rupp et al., 2025; Santoro et al., 2014, 2017). However, the computational advantage of such an organization has remained unclear. The present results suggest that the principal computational advantage of this organization emerges under causal listening conditions. When future acoustic information is unavailable, parallel processing across multiple temporal contexts reduces the ambiguity associated with incomplete sensory evidence and substantially improves sound recognition. In this sense, multiscale processing provides a computational strategy for progressively refining perceptual interpretations as acoustic evidence accumulates over time (Honey et al., 2012).

The comparison between causal and non-causal architectures is important because it argues against a purely capacity-based explanation of the multistream advantage. Although adding streams increases the dimensionality and expressive capacity of the model, such an explanation would predict comparable benefits in causal and non-causal settings. Instead, the advantage is markedly larger when the model must recognize sounds without access to future acoustic information. This finding suggests that the parallel streams provide complementary temporal descriptions that become particularly valuable in the absence of bidirectional context. In this sense, multiscale processing compensates for a computational limitation specific to real-time perception.

Viewed more broadly, the present results suggest that multiscale temporal processing is not merely a descriptive property of biological audition, but an adaptive computational strategy for reducing uncertainty during causal inference from temporally evolving sensory signals. The proposed architecture should therefore be viewed as a brain-inspired computational account of causal auditory recognition, rather than as a literal model of cortical circuitry.

### 3.0.2 How Multiscale Processing Shapes Internal Representations

The temporal-modulation reconstruction analysis provides a mechanistic account of how information is represented and transformed across the recurrent hierarchy. Temporal modulations are fundamental components of auditory representations and have been used extensively to characterise multiresolution sound processing in auditory systems (Chi et al., 2005; Santoro et al., 2014, 2017). In the first recurrent layer, all three temporal streams retain substantial information about the modulation structure of the sounds. The 80 ms stream exhibits the broadest and strongest reconstruction, whereas the combined representation closely resembles the longest stream. This pattern suggests that the first recurrent stage primarily preserves stream-dependent acoustic information and that much of the recoverable modulation structure is already captured by the longest temporal context.

A different organisation emerges in the second recurrent layer. Reconstruction from individual streams becomes less extensive, particularly for intermediate and higher modulation rates, indicating a progressive transformation away from a detailed acoustic description of the input. At the same time, the combined representation retains a broader and more stable range of temporal-modulation information than any single stream alone. This pattern is consistent with the integration of complementary information across temporal scales. The network therefore does not merely accumulate acoustic information across layers. Instead, recurrent processing appears to reorganise stream-dependent representations into a more integrated multiscale code. This code retains selected temporal information while discarding part of the fine-grained acoustic structure present at earlier stages.

Although these analyses do not formally demonstrate synergistic information across streams (Williams and Beer, 2010), they show that the combined representation becomes more informative than any individual stream at the deeper recurrent stage. This suggests that the computational advantage of multiscale processing emerges through hierarchical integration rather than from a single dominant temporal stream.

The RSA results provide convergent evidence for this account at the level of categorical separability (Kriegeskorte et al., 2008). Early convolutional layers show weak correlation with the categorical model across all temporal streams, indicating that local spectro-temporal filtering alone does not produce strongly organised category representations. A first increase in correlation with the categorical model appears in the initial recurrent layer, with peaks around 1.5–2 s, followed by a slower and more sustained increase in the deeper recurrent layer. Importantly, the isolated branches of the multiscale network are not consistently stronger than their matched single-stream counterparts. In several recurrent-layer comparisons, the single-stream models reach comparable or higher categorical separability within the corresponding temporal branch. The clearest multiscale advantage appears only after stream concatenation and classification, where category-level organisation becomes substantially stronger than in any individual stream. This pattern indicates that the main representational benefit of the multiscale architecture lies in fusion and readout rather than in a uniform strengthening of every temporal branch.

Taken together, the temporal-modulation and RSA analyses suggest a coherent hierarchical transformation. Early processing stages preserve distinct aspects of the incoming acoustic signal across temporal windows. Recurrent processing subsequently transforms and combines these representations, producing a shared multiscale representation that supports stronger category-level organisation after stream fusion. This interpretation is consistent with accounts of auditory perception that emphasise temporal context, recurrent processing, and hierarchical integration in the formation of stable auditory objects (Elhilali et al., 2009; Poeppel et al., 2012; Bizley and Cohen, 2013; Giordano et al., 2023). The present findings extend these accounts by identifying a specific computational role for multiscale temporal integration under causal constraints.

#### Bioinspired Architectures for Hypothesis Generation

An important contribution of this work is methodological. Because the proposed model operates causally and produces framewise internal representations, its activity can be aligned directly with time-resolved neural measurements such as EEG, MEG, and intracranial electrophysiology. This creates a practical framework for testing whether biological auditory systems exhibit a comparable progression from stream-dependent acoustic representations toward integrated multiscale codes. More specifically, the model generates testable predictions about when temporal-modulation information is preserved, when representations from different temporal scales are combined, and when category-level organisation becomes most strongly expressed. These properties make the MSCRNN architecture a useful hypothesis-generating framework for studying the temporal dynamics of auditory recognition.

This idea is consistent with a broader trend toward neuroanatomically informed deep learning models. Explicit implementations of neuroanatomic structure into deep neural network models leads to biologically more plausible neurocomputational mechanisms in vision (e.g., Weidler et al., 2021; Lu et al., 2025) and motor control (Weidler, 2025). Beyond the inductive bias on the emergence of the mechanism, neuroanatomical structure facilitates a direct mapping between *in vivo* and *in silico* components. Through this mapping, revealed mechanisms in the model lead to possible explanations of cortical processes (Weidler, 2025). In the same way, the present work proposes a model architecture that allows us to explicitly probe anatomically distinct pathways (here streams) for their representational and transformational characteristics.

#### Limitations and future extensions

Although the present results establish a computational role for multiscale processing under causal constraints, several limitations should be considered. First, the model was evaluated on isolated environmental sounds rather than complex auditory scenes involving overlapping sources. Consequently, the selected 20–40–80 ms temporal contexts may not generalise to more challenging listening conditions. Second, the three-stream architecture represents only a simplified approximation of biological temporal heterogeneity. Auditory cortex is unlikely to operate with only three discrete temporal contexts, and the optimal temporal organization n may vary across cortical fields and auditory tasks (Chi et al., 2005). Third, F1 evaluation, temporal-modulation reconstruction and RSA analyses are correlational and do not establish the causal contribution of individual streams or layers. Targeted ablations, variance partitioning, and partial-information decomposition will be required to distinguish unique, redundant, and synergistic contributions across temporal scales (Seibold and McPhee, 1979; Williams and Beer, 2010). Fourth, the RSA analyses rely on discrete sound categories and may overlook more continuous representational dimensions. Finally, the hyperparameter search was not exhaustive, and should not be interpreted as identifying a globally optimal architecture. Rather, it demonstrates that simply increasing temporal context and architectural complexity does not necessarily improve performance. The present architecture should therefore be regarded as one effective brain-inspired solution rather than a definitive model of auditory cortical organisation.

A particularly promising next step will be to compare the internal representations of alternative MSCRNN architectures directly with time-resolved neural activity. Such comparisons can determine whether architectures optimised for sound recognition also provide the best account of cortical processing, or whether biologically motivated temporal organisations more closely match the dynamics of auditory representations measured with techniques such as magneto-encephalography(MEG). In this way, the proposed framework enables optimisation with respect to both behavioural performance and alignment with neural data.

Beyond neural validation, future work should extend this framework to naturalistic scenes with overlapping sources, and should not be interpreted as identifying a globally optimal architecture. Rather, it demonstrates that simply increasing temporal context and architectural complexity does not necessarily improve performance. The present architecture should therefore be regarded as one effective brain-inspired solution rather than a definitive model of auditory cortical organisation. Continuous semantic embeddings (Esposito et al., 2024) may provide a richer description of representational organisation than discrete sound categories. Taken together, the present results identify multiscale temporal processing as a plausible computational strategy for robust auditory recognition when future acoustic information is unavailable.

## 4 Conclusion

This work identifies multiscale temporal processing as a computational strategy for auditory recognition under causal constraints. When future acoustic information is unavailable, processing sounds across multiple temporal contexts substantially reduces the performance cost of online recognition, allowing causal models to approach the accuracy of non-causal architectures after an initial period of evidence accumulation. This advantage cannot be explained simply by increased model capacity, but reflects the particular value of parallel temporal processing when recognition must unfold in real time.

Mechanistically, our analyses suggest that multiscale processing supports a hierarchical transformation of sound representations. Early recurrent processing preserves complementary acoustic information across temporal contexts, whereas deeper processing combines these representations into increasingly category-related representations while retaining behaviourally relevant temporal dynamics.

Together, these findings provide a computational rationale for why biological auditory systems may analyse sounds over multiple temporal contexts. Beyond improving causal sound recognition, the proposed MSCRNN architecture offers an interpretable framework for generating and testing hypotheses about the temporal organisation of auditory processing through direct comparison with time-resolved neural recordings.

## Declarations

## Acknowledgments

This work was supported by: the European Union’s Horizon Europe research and innovation programme under the Marie Skl-odowska-Curie Actions (Grant Agreement No. 101279402, SONATA), the European Research Council(ERC-2024-SyG NASCE, Grant Agreement No. 101167313), the French National Research Agency (ANR-21-CE37-0027-01; ANR-16-CONV-0002 ILCB; ANR-11-LABX-0036 BLRI), and the Dutch Research Council (NWO Grant No. 406.20.GO.030). This work used the Dutch national e-infrastructure with the support of the SURF Cooperative (Grant No. EINF-14563), the Data Science Research Infrastructure (DSRI; Maastricht University), and the Province of Limburg.

## Data Availability

The code used to train, evaluate, and analyse the MSCRNN models will be made publicly available on GitHub upon publication. The SHDC pre-training dataset was constructed from audio recordings licensed from Sound Ideas Inc. (Sound Ideas Inc., 2022b,a) and therefore cannot be redistributed by the authors. Access to the original recordings must be obtained directly from Sound Ideas under the appropriate licence. The ESC-50 dataset (Piczak, 2015) is publicly available. The trained model weights will also be made available through the same GitHub repository upon publication.

## Methods

### 4.1 Model architecture

The model architecture is illustrated in Figure 1. The MSCRNN consists of three parallel processing streams operating over distinct temporal windows. At every 10 ms time step, the log-Mel spectrogram is extended by one newly computed frame. Three sliding temporal windows of different durations are then extracted from the most recent portion of the spectrogram. Each stream independently processes its input using a convolutional layer, a spectral attention layer, and two recurrent layers. The resulting stream-specific representations are concatenated and passed through a softmax layer that predicts framewise probabilities over classes. Unless otherwise stated, all models operate causally, such that only present and past acoustic information is available at each time step.

#### 4.1.1 Temporal scales and time constants

Each stream receives acoustic information through a sliding time window. Two temporal scale regimes were evaluated. In a configuration motivated by the neural analysis of speech-processing, scales of 50, 200, and 400 ms were used, reflecting temporal integration timescales reported in human auditory cortex (Norman-Haignere et al., 2022). In a task-optimised configuration, 20, 40, and 80 ms were selected via hyperparameter optimisation on a subset of the pre-training dataset (Supplementary Methods Section S-M4). At each time step, the window only contains present and past log-Mel spectrogram frames, ensuring causality in the model’s access to the unfolding acoustic input.

#### 4.1.2 Convolutional layers

Within each stream, convolutional layers extract local spectro-temporal features from the time window available at every time step. Because the temporal scales differ in duration, different convolutional kernel geometries were used across streams. Following sequential architecture calibration, the selected configuration employed kernels of 1 × 5, 5 × 5, and 5 × 1 for the short-, intermediate-, and long-duration streams, respectively (Supplementary Methods Section S-M4; Supplementary Tables S1-S2). These asymmetric kernels provide the shorter scales with greater opportunity to learn fine temporal structure while allowing longer scales to capture broader spectral patterns over their larger temporal context.

#### 4.1.3 Attention mechanisms

The inclusion of attention mechanisms were tested at the output of both convolutional and recurrent layers to enhance the weighting of informative spectral and temporal features. In convolutional layers, attention operates only along the frequency dimension, whereas in recurrent layers it selectively weights time steps according to their contribution to sound categorisation. The inclusion and placement of attention modules were treated as hyperparameters; optimization results showed that a single attention layer at the output of the convolutional layer provided the best performance (Figure 1). Implementation details are provided in Supplementary Methods Section S-M2, while the tested configurations and optimisation results are reported in Supplementary Table S5 and Supplementary Figure S5.

#### 4.1.4 Recurrent layers and directionality

Following convolution and spectral attention, the feature sequence produced by each temporal stream is processed by two stacked gated recurrent unit (GRU) layers comprising 512 and 128 hidden units, respectively. The recurrent layers integrate convolutional features over time, enabling the network to accumulate information beyond the local temporal receptive field of the convolutional filters. Two recurrent configurations were evaluated. In the *causal* configuration, both recurrent layers consist of forward GRUs, such that the representation at time *t* depends exclusively on the current input and all preceding time steps. Consequently, the prediction associated with each time point can be computed without access to future acoustic information. In the non-causal configuration, the forward GRUs are replaced by bidirectional GRUs. During both training and inference, the complete sequence of convolutional features is processed in both forward and backward directions before the hidden representations are computed. Consequently, the representation at time *t* incorporates information from both preceding and subsequent frames of the input sequence. The non-causal architecture therefore provides an offline upper bound against which the performance of the causally constrained model can be evaluated. Note that, although the non-causal models have access to the complete acoustic sequence through bidirectional recurrence, predictions at each time point view the sequence divided differently over forward and backward pass. Therefore, temporal blurring effects shift from mostly affecting late to mostly affecting earlier time steps. Framewise performance thus varies over time, reflecting differences in the perspective on the sequence at different portions of the sound rather than limitations imposed by causal processing (Supplementary Methods Section S-M4, with the final configurations reported in Supplementary Tables S5–S6).

### 4.2 Pre-processing

All audio signals were resampled to 16 kHz, converted to mono, and normalised to a duration of 5 s. Sounds shorter than 5 s were looped to match this duration. Log-Mel spectrograms were computed using a short-time Fourier transform with a 25 ms Hanning window and a hop size of 10 ms, resulting in 128 Mel-frequency bands spanning 125–7500 Hz. At each 10 ms time step, each stream extracted a causal sliding temporal window spanning the current and preceding acoustic context. Windows extending beyond the beginning of the sound were left-padded with zeros, ensuring that no future information was available to the model. For a 5 s input, this yielded windowed spectrograms of dimensions 498*×*5*×*128, 498*×*20*×*128, and 498*×*40*×*128 for the 50 ms, 200 ms, and 400 ms temporal windows, respectively. Silent frames were identified using an RMS-energy threshold and handled using the zero-label strategy selected during architecture calibration. Full details are provided in Supplementary Methods Section S-M3. The distribution of the dataset utilised for this work (see below) is shown in Supplementary Figure S1, while the comparison of alternative frame-labelling strategies is reported in Supplementary Figure S7 and Supplementary Table S7.

### 4.3 Dataset

The MSCRNN was trained in a two-stage procedure. Model weights were first pre-trained on a large-scale, weakly structured audio corpus and subsequently fine-tuned on a curated environmental sound classification dataset.

#### 4.3.1 Pre-training dataset

Pre-training was performed using a subset of the *SuperHardDrive Combo (SHDC)* dataset (Sound Ideas Inc., 2022b), which contains 388,199 variable-length audio recordings spanning a broad range of sound sources, including speech, environmental sounds, animal vocalisations, and mechanical noises. Although SHDC labels are loosely defined and heterogeneous, the dataset provides substantial acoustic diversity, making it well suited for learning general-purpose spectro-temporal feature representations.

To reduce semantic redundancy and class imbalance, we constructed a semantically balanced subset following the procedure described by Esposito et al. (2024). Briefly, Word2Vec embeddings computed from the textual sound labels were hierarchically clustered, and clusters were ordered according to their inter-cluster cosine distance to maximise semantic diversity. From an initial set of 300 clusters, the 260 most semantically distinct clusters were retained. Each cluster contributed at most 2,000 sound files to the pre-training corpus, whereas smaller clusters were included in full. Considering the textual labels of the sounds retained in any of the cluster, resulted in 612 distinct classes, which were one-hot encoded.

To match the fixed 5 s input requirement of the MSCRNN, recordings longer than 5 s were segmented into multiple non-overlapping excerpts. Each excerpt was treated as an independent training example, increasing the effective size of the pre-training corpus to 1,210,329 five-second sound segments while preserving the approximately balanced semantic distribution established during the initial sampling procedure. Further details of the semantic balancing and dataset expansion are provided in Supplementary Methods Section S-M1.

### 4.4 Fine-tuning dataset

After pre-training, the model was fine-tuned on the *ESC-50* dataset (Piczak, 2015), a benchmark dataset specifically designed for environmental sound classification. ESC-50 consists of 2,000 five-second recordings distributed across 50 balanced classes grouped into animal, natural, and human-related sound categories. The dataset is organised into five predefined folds of 400 sounds each. Fine-tuning was performed on the first four folds, while the evaluation results reported in this study were based on the fifth fold, which served as the held-out test set.

### 4.5 Training Procedure

Training the MSCRNN is computationally demanding because each architecture processes long frame sequences through three parallel convolutional-recurrent streams. To make architecture calibration computationally feasible, hyperparameter optimisation was performed on a reduced subset of 23,255 sounds sampled from the SHDC dataset. Once the best-performing architecture had been identified, it was retrained from scratch on the complete SHDC pre-training corpus before being fine-tuned on the ESC-50 dataset and evaluated on the held-out fifth fold.

Model parameters were optimised using categorical cross-entropy loss,

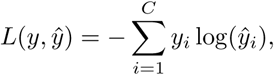

where *C* denotes the number of classes, *y_i_*the ground-truth label, and *y*^*_i_* the predicted class probability. Optimisation was performed using Adam (Kingma and Ba, 2014) with an initial learning rate of 10*^−^*^3^ that decayed exponentially with a rate constant of 0.1.

Before each epoch, training sounds were randomly shuffled to avoid sequential biases and to ensure that each 5 s excerpt was treated as an independent training example. Silent frames were identified using an RMS-energy threshold and assigned a dedicated zero label, which was selected during architecture calibration as the best-performing silence-handling strategy (Supplementary Methods Section S-M3).

During fine-tuning, all layers up to and including GRU1 were frozen, while only the final classification layer was updated. This freezing depth was selected empirically by comparing alternative fine-tuning strategies (Supplementary Figure S9). All training was performed on NVIDIA A100 GPUs with 40 GB of memory.

Full details of the sequential calibration procedure are provided in Supplementary Methods Section S-M4. The explored search spaces and selected architectures are summarised in Supplementary Tables S1–S7, and intermediate optimisation results are reported in Supplementary Figures S2–S7, including the temporal-window search (Supplementary Figure S6; Supplementary Table S6).

### 4.6 Evaluation Metrics

Model performance was evaluated using the framewise F1-score (Mesaros et al., 2016). This metric is particularly suited to time-resolved sound event classification, as it assesses prediction accuracy independently at each temporal frame.

#### 4.6.1 Framewise F1-Score

The framewise F1-score is defined as the harmonic mean of precision and recall, computed at the level of individual time frames:

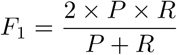

where precision *P* and recall *R* are given by:

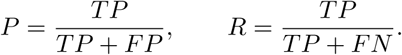

Here, *TP*, *FP*, and *FN* denote the number of true positives, false positives, and false negatives, respectively, computed per frame.

#### 4.6.2 Confidence Interval Estimation and Significance Testing

From 10 trainings per model type on independent seeds we estimate the expected value of the model performance (F1-score) time-point-wise. Confidence intervals are calculated from a bias-corrected and accelerated (BCa) bootstrap (Efron, 1987) with runs independently drawn *B* = 10 000 times within conditions, with a nominal coverage of 95%.

To test the pointwise significance of the difference of differences *δ* reported in Figure 2c, we invert percentile (not BCa) confidence intervals into p-values and determine the fraction of bootstrap resamples below zero (*H*_0_: *δ* = 0). To control for *T* = 498 repeated tests (time points) with positive dependence induced by temporal autocorrelation, we apply a false discovery rate (FDR) correction under the Benjamini-Hochberg procedure (Benjamini and Hochberg, 1995) at *α* = 0.05.

### 4.7 Representational Similarity Analysis

To characterise the emergence of task-relevant categorical information throughout the MSCRNN hierarchy, we performed a time-resolved Representational Similarity Analysis (RSA) (Kriegeskorte et al., 2008). Rather than relying on the network’s final classification output, this analysis quantifies the extent to which the representational geometry within each layer reflects the target sound categories. Consequently, RSA provides a classifier-independent measure of how well the internal representations separate the ESC-50 classes across processing stages and over time.

For each layer and each time step, we computed a representational dissimilarity matrix (RDM) across sounds using correlation distance:

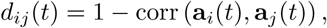

where **a***_i_*(*t*) denotes the activation pattern elicited by sound *i* at time *t*. Correlation distance captures differences in representational structure while remaining largely insensitive to global activation magnitude, facilitating comparisons across layers with different activation scales and nonlinearities.

Each RDM summarises the pairwise similarity structure of the sound representations at a given time point, providing a time-resolved description of how internal representations evolve throughout sound processing.

To quantify the emergence of category-selective representations, we constructed a model RDM based on the ground-truth ESC-50 labels. The categorical model encoded the 50 individual sound classes by assigning a low dissimilarity to sounds belonging to the same category and a high dissimilarity to sounds belonging to different categories. The high correspondence between a layer RDM and this model therefore indicates that sounds from the same category have become increasingly separated from sounds belonging to different categories within that layer’s representational space.

#### 4.7.1 RSA Correlation Timecourses

For each layer, the representational alignment with the categorical model was quantified by computing the Pearson correlation between the vectorized layer RDM and the categorical model RDM at every time step.

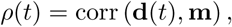

where **d**(*t*) denotes the vectorised layer RDM at time *t*, and **m** denotes the vectorised categorical model RDM.

This procedure yields a time-resolved profile describing when task-relevant category information becomes most strongly expressed within each layer. For each layer, we identified the time of peak correlation as the point at which category separability was maximal.

### 4.8 Temporal modulation reconstruction analysis

To investigate how temporal information is represented across the network hierarchy, we quantified the relationship between the internal representations of the causal MSCRNN and the temporal modulation structure of the input sounds (Chi et al., 2005).

#### 4.8.1 Causal temporal modulation representation of sounds

Temporal modulation representations were estimated from the sound waveforms (ESC-50 dataset) using a causal auditory-inspired modulation analysis pipeline (Lindeberg and Friberg, 2015). Spectral magnitudes were compressed using logarithmic compression and averaged across frequency channels to derive a broadband auditory envelope trajectory sampled with 10 ms temporal resolution. Temporal modulation trajectories were then estimated by filtering the envelope with a bank of strictly causal Butterworth temporal modulation filters (order=2) tuned to modulation rates between 0.5 and 96 Hz, logarithmically spaced across 32 channels. The slowest channel was implemented as a low-pass filter, while higher modulation rates were estimated using causal band-pass filters with bandwidth controlled by a quality-factor parameter (Q=1). The absolute value of the filter outputs was retained to obtain instantaneous modulation-energy trajectories over time. This causal implementation ensured that temporal modulation estimates at each time point depended only on present and past acoustic information, matching the causal architecture of the CRNN model. Temporal modulation representations were computed continuously over the 5 s sound duration and subsequently averaged within selected temporal windows for representational analyses.

#### 4.8.2 Regularized canonical correlation analysis

To characterize the temporal modulation information represented in the recurrent network layers, we used regularized canonical correlation analysis (regCCA, Vinod (1976); Leurgans et al. (1993); Uurtio et al. (2018)) to reconstruct temporal modulation features from recurrent activations. The analysis was performed in a sliding-window manner using successive 200 ms temporal windows advanced in 100 ms steps, resulting in 50 overlapping analysis windows spanning the duration of each sound. For each window, the acoustic representation consisted of the temporal modulation spectrum of each sound (32 temporal modulation frequency bins) averaged over the corresponding 200 ms interval. Network representations were obtained by computing, for every recurrent unit of GRU0 or GRU1, the Euclidean norm of its activity across the same 200-ms window. Before windowing, the temporal mean of each unit’s activation was removed across the sound. For the multistream model, activation vectors from the 20-, 40-, and 80-ms streams were first normalized separately to unit global root-mean-square energy and then concatenated into a single feature vector. This normalization prevented streams with larger activation magnitudes from dominating the multivariate analysis. For single-stream analyses, the same procedure was applied to the activation vector from the selected stream only. Thus, the subsequent regCCA and dimensionality-reduction procedures were identical for the one- and three-stream analyses; the only difference was whether the input feature matrix contained one stream or the concatenation of all three streams.

### 4.9 Nested cross-validation and dimensionality reduction

Reconstruction performance was evaluated using nested cross-validation. The outer loop consisted of five folds, such that 80% of the sounds formed a development set and the remaining 20% formed an independent test set. Each sound therefore received exactly one prediction from a model for which that sound had not contributed to preprocessing, dimensionality reduction, hyperparameter selection, or model fitting. Within each outer development set, five-fold inner cross-validation was used to jointly select the regularization parameters for the acoustic and network representations, *λ_x_* and *λ_y_*, and the number of canonical components used for reconstruction. Both regularization parameters were searched over eight logarithmically spaced values (*λ ∈* [10*^−^*^3^, 10^4^]) and the number of canonical components was varied from one to five. Because the recurrent representations contained substantially more features than sounds, principal component analysis (PCA) was applied to the standardized network activations before regCCA. PCA was fitted exclusively on the network data from the relevant training partition and retained 100 components. The fitted PCA transformation was then applied without refitting to the corresponding validation or test data. In the three-stream analysis, PCA was performed jointly on the concatenated activation vector from all three streams. Consequently, the resulting principal components could express variance shared across streams as well as variance specific to an individual stream. No separate PCA was performed within each stream. In the single-stream analyses, PCA was instead fitted to the activation matrix of the selected stream alone, using the same 100 components and the same fold-specific fitting procedure. For every candidate combination of *λ_x_*, *λ_y_*, and canonical dimensionality, a reconstruction model was fitted to the inner-training partition and used to predict the acoustic modulation features of the inner-validation sounds. Reconstruction accuracy was quantified separately for each temporal modulation rate using the Pearson correlation between observed and predicted values across validation sounds. Correlations were Fisher-z transformed and averaged across modulation rates and validation folds. The hyperparameter combination yielding the highest mean validation score was selected independently within each outer fold.

#### 4.9.1 RegCCA fitting and reconstruction

Following hyperparameter selection, preprocessing, PCA, regCCA, and the reconstruction mapping were refitted using the complete outer development set. RegCCA was estimated from the regularized covariance matrices

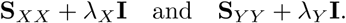

where *X* denotes the temporal modulation representation and *Y* the PCA-reduced recurrent representation. Singular-value decomposition of the whitened cross-covariance matrix yielded canonical projection matrices for the two representational spaces.

The recurrent activations were projected into the selected canonical space, and a linear reconstruction mapping from the recurrent canonical scores to the standardized temporal modulation features was estimated using the Moore–Penrose pseudoinverse. This mapping was then applied to the recurrent representations of the held-out outer-test sounds. Predicted modulation features were finally transformed back to their original feature scale. Predictions from all five outer folds were concatenated, and reconstruction performance was quantified for each temporal modulation rate using both Pearson correlation between the observed and independently predicted values across all sounds.

#### 4.9.2 Permutation testing

Statistical significance was assessed using 1,000 permutations while retaining the outer-fold assignments and the hyperparameters selected from the non-permuted data. Within each outer fold and permutation, the correspondence between the temporal modulation features and recurrent representations was randomly shuffled only within the outer development set. Specifically, the rows of the development-set recurrent representation were permuted relative to the acoustic features. The complete preprocessing, PCA, regCCA, and reconstruction mapping were then refitted using this permuted development correspondence. The outer test data were not permuted. Each null model was therefore evaluated using the original, correctly paired recurrent and acoustic representations of the held-out test sounds. This procedure tests whether a model trained without a systematic correspondence between acoustic modulation features and network representations can nevertheless reconstruct the correctly paired unseen sounds.

For every permutation, the independently predicted outer-test values were combined across folds and feature-wise Pearson *r* values were calculated. For each modulation frequency and time window, the observed reconstruction accuracy was normalized by subtracting the mean of the null distribution and dividing by its standard deviation, yielding a z-scored reconstruction measure that quantified encoding strength relative to chance. Modulation ranges that were represented above chance level (*Z >* 3, *p <* 0.001, one-tailed, uncorrected) are visualized in yellow in the heat map.

## Supplementary Material

Multiscale Temporal Processing Supports Sound Recognition under Causal Constraints

## 1 Supplementary Methods

This Supplementary Material provides technical details and extended analyses supporting the main manuscript. The first section describes dataset preparation, attention implementation, silence handling, sequential hyperparameter optimisation, representational similarity analysis, and temporal-modulation reconstruction. The second section reports the optimisation results used to calibrate the MsCRNN architecture, including the selection of the 20–40–80 ms temporal-buffer regime, as well as robustness analyses of silence handling and fine-tuning depth. The final section presents extended representational analyses comparing the task-optimised temporal regime with the longer 50–200–400 ms configuration motivated by prior auditory-neuroscience research.

### 1.1 S-M1. Dataset expansion and semantic balancing

To mitigate class imbalance and semantic redundancy in the SuperHardDrive Combo (SHDC) dataset, sounds were sampled from a semantically balanced subset generated through hierarchical clustering using word2vec embeddings [Mikolov et al., 2013] derived from textual sound labels [Esposito et al., 2024]. Clusters were ranked according to inter-cluster cosine distance to promote semantic diversity. From an initial set of 300 clusters, the 260 most semantically distinct clusters were retained. Each cluster contributed up to 2,000 sound files, while clusters containing fewer samples were included in full.

Long audio recordings were segmented into non-overlapping 5 s excerpts to match the fixed-duration model input, for a total of 1,210,329 five-second segments in the final pre-training corpus (*≈*1,681 hours of audio).

### 1.2 S-M2. Frequency-wise attention implementation

To apply attention along the spectral dimension, convolutional feature maps were permuted to expose the frequency axis explicitly. Given an input tensor of shape (*T, B, F, C*), the tensor was permuted to (*T, F, B, C*) prior to attention computation.

Channel-averaged activations were used to compute frequency attention scores:

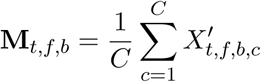

These scores were normalised using a softmax function and applied multiplicatively to the original feature maps before reverting to the original tensor ordering.

### 1.3 S-M3. Temporal Attention in GRU Layers

Temporal attention was applied to GRU outputs to emphasise informative time steps. Given GRU activations **H** *∈* R*^T^ ^×F^*, attention weights were computed as:

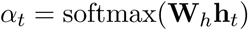

The resulting context vector was obtained as a weighted sum:

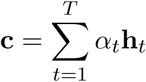

This mechanism allows the network to adaptively prioritise temporal regions relevant for sound classification.

Alternative attention placements were evaluated during hyperparameter optimisation, including attention after recurrent layers and combinations of spectral and temporal attention modules. The full set of tested conditions is reported in Table S5 and Figure S5.

### 1.4 S-M3. Silence detection and frame-labelling strategies

Because the MSCRNN produces framewise predictions at a temporal resolution of 10 ms, each time step contributes independently to the training loss. However, many frames correspond to silence or contain only low-energy acoustic information, providing little category-discriminative information. We therefore evaluated alternative strategies for handling these frames during training.

Silence was identified using the root mean square (RMS) energy of the spectrogram at each time step:

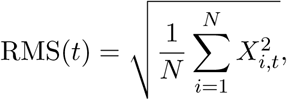

where *X_i,t_*denotes normalized spectrogram power at frequency bin *i* and time *t*, and *N* = 128 is the number of Mel-frequency bins. Frames with RMS energy below a fixed threshold of 0.1 were classified as silent.

Figure S1 shows the temporal distribution of silent frames and the distribution of silence percentages across sounds in SHDC and ESC-50.

Three frame-labelling strategies were compared. Under *label repetition*, the global sound label was assigned to every frame, including silent frames. Under the *silence-class strategy*, low-energy frames were assigned to a dedicated silence category. Under the *zero-label strategy*, all elements of the target vector were set to zero for silent frames. These frames therefore contributed no class-specific cross-entropy term while remaining included in the reduction of the loss across time steps.

The zero-label strategy yielded the highest validation performance during architecture calibration and was used in subsequent analyses. The complete comparison is shown in Figure S7 and Table S7.

### 1.5 S-M4. Sequential hyperparameter optimisation

We calibrated the MsCRNN architecture before evaluating the consequences of strict causal processing. The search began from a biologically motivated multiscale prior based on temporal integration windows reported in auditory cortex during speech-related processing (50–200–400 ms; Norman-Haignere et al., 2022). Architectural parameters were then varied sequentially to identify a configuration better matched to the statistical properties of environmental sounds.

**Table S1:** Summary of the hyperparameter optimisation space and selected MsCRNN configuration. The complete search space is reported in Tables S2–S7.

| Model component | Explored configurations | Selected configuration |
| --- | --- | --- |
| Convolutional filter geometry | Asymmetric time–frequency kernels adapted to buffer duration; square and reversed control kernels (Table S2) | Short: $1 \times 5$<br>Intermediate: $5 \times 5$<br>Long: $5 \times 1$ |
| Convolutional depth and filter count | One or two convolutional layers with 4–16 filters per layer (Table S3) | Single convolutional layer with 4 filters |
| Recurrent architecture | GRU layers with 128–1024 units; single-layer and multi-layer variants (Table S4) | Two-layer GRU (512, 128 units) |
| Attention placement | No attention; convolutional attention; recurrent attention; combined attention variants (Table S5) | Attention after first convolutional layer |
| Temporal-buffer sizes | Short-scale, logarithmic, neuroscientifically motivated, and equal-window control configurations (Table S6) | 20–40–80 ms |
| Silence handling | Label repetition; dedicated silence class; zero-label strategy (Table S7) | Zero-label strategy |

Hyperparameter optimisation was conducted using the less restrictive recurrent configuration, in which bidirectional recurrence allowed the network to exploit the information available within each temporal buffer. This design choice served a specific methodological purpose: it allowed temporal-scale discovery without prematurely constraining the search through the additional computational demands imposed by strictly forward recurrence. After selecting the task-optimised configuration, including the 20–40–80 ms temporal regime, we re-evaluated the resulting architecture under strict causal processing in the main analyses.

Given the scale of the pre-training corpus, optimisation was performed on a reduced subset of 23,255 sounds sampled from SHDC. Configurations were compared using framewise F1 score on held-out validation data. The search proceeded sequentially across convolutional kernel geometry, convolutional depth and filter count, recurrent depth and dimensionality, attention placement, temporal-buffer configuration, and silence-handling strategy. Because framewise training produces many silent or low-energy frames that provide little category-discriminative information, alternative strategies for assigning labels to these frames were also evaluated during optimisation. This sequential procedure reduced computational cost while preserving a consistent architecture across temporal streams. The selected model was subsequently retrained on the full SHDC corpus and fine-tuned on ESC-50 before all subsequent analyses.

Table S1 summarizes the complete optimisation sequence and the final selected architecture. Detailed search spaces are reported in Tables S2–S7.

### 1.6 S-M5. Representational similarity analysis

Representational similarity analysis (RSA) was used to characterize how category-level structure evolves across network layers and time [Kriegeskorte et al., 2008]. Activations were extracted from all layers of interest for the 400 held-out ESC-50 sounds in fold 5.

For each layer and time point, each sound was represented by a feature vector containing the corresponding hidden activation pattern. A representational dissimilarity matrix (RDM) was then computed across sounds using correlation distance:

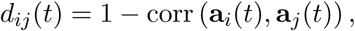

where **a***_i_*(*t*) denotes the activation vector elicited by sound *i* at time *t*.

A categorical model RDM was constructed from the 50 ESC-50 class labels. Stimulus pairs belonging to the same category were assigned dissimilarity 0, while pairs belonging to different categories were assigned dissimilarity 1. For each layer and time point, the upper triangular portions of the empirical and model RDMs were vectorized and correlated using Pearson correlation:

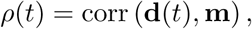

where **d**(*t*) denotes the vectorized empirical RDM and **m** denotes the categorical-model vector. The resulting RSA time courses quantify the extent to which internal model representations reflect category-level structure as the sound unfolds.

## 2 Supplementary Results

### 2.1 S-R1. Sequential architecture optimisation

The sequential optimisation procedure was used to identify an architecture suitable for the subsequent causal analyses. All model configurations were evaluated using framewise F1 score at a temporal resolution of 10 ms.

Figure S2 shows the search over convolutional kernel geometries. Several configurations yielded comparable performance. The selected configuration used kernels of 1 *×* 5, 5 *×* 5, and 5 *×* 1 for the short, intermediate, and long streams, respectively.

Figure S3 reports the optimisation of convolutional depth and filter count. The best-performing configuration consisted of a single convolutional layer with four filters.

Figure S4 reports the optimisation of recurrent depth and dimensionality. A two-layer GRU architecture with 512 and 128 units yielded the highest validation performance.

Figure S5 shows the comparison of attention placements. Applying attention immediately after the first convolutional layer yielded the highest validation performance.

Figure S6 shows the comparison of temporal-buffer configurations. The [2, 4, 8] configuration, corresponding to temporal windows of 20, 40, and 80 ms, achieved the highest validation framewise F1 score.

Figure S7 shows the comparison of silence-handling strategies. The zero-label strategy yielded the highest validation framewise F1 score and was retained for subsequent analyses.

### 2.2 S-R2. Robustness to silence handling

To determine whether the principal causal and multiscale comparisons depended on the explicit treatment of silent frames, we compared the two configurations once more, without applying the selected zero-label strategy. Figure S8 shows the resulting time-resolved framewise F1 scores.

The qualitative pattern remained consistent with the main analyses. Multistream processing provided a substantial advantage in the causal condition, whereas differences between single-stream and multistream models were smaller in the non-causal condition. This analysis indicates that the central interaction between causal processing and multiscale integration is not driven exclusively by the treatment of silent frames.

### 2.3 S-R3. Effect of freezing depth during ESC-50 fine-tuning

We evaluated the effect of freezing depth during ESC-50 fine-tuning using the task-optimised 20–40–80 ms MsCRNN architecture. Alternative strategies included freezing all feature-extraction layers up to the classifier, freezing weights up to the second recurrent layer (GRU1), freezing weights up to the first recurrent layer (GRU0), and fine-tuning the complete network.

Performance depended on the selected freezing depth. The highest framewise F1 scores were obtained when weights were frozen up to GRU1, whereas full-network fine-tuning yielded lower performance. This pattern suggests that representations acquired during large-scale SHDC pre-training remain transferable to ESC-50 and that excessive adaptation can degrade useful spectro-temporal representations.

### 2.4 S-R4. Additional multi- and single-scale architectures

To investigate whether larger temporal context windows in the causal multiscale architecture still outperform their constituents, we also test a multiscale architecture with a 50–200–400ms regime and compare it to its 50ms, 200ms, and 400ms single-stream variants Figure S10. As for the 20–40– 80ms regime, the 50–200–400ms regime significantly outperformed the single-stream constituents. Additionally, Figure S10 shows that larger context sizes (and hence more parameters) does not improve the performance of the multistream design.

### 2.5 S-R5. Number of model parameters

Figure S11 shows the number of trainable parameters in each individual model variant of multi- and single-stream designs. Naturally, multiscale models count more parameters than their individual constituents.

### 2.6 S-R6. Extended RSA comparison across temporal-integration regimes

To determine whether category-level representational dynamics depend on the choice of temporal-integration windows, we repeated the RSA analysis using the 50–200–400 ms comparison architecture motivated by prior work on speech-related temporal integration in auditory cortex [Norman-Haignere et al., 2022].

Figure S12 shows the corresponding category-level RSA time courses. At early processing stages, the two architectures exhibit broadly similar dynamics. In the first recurrent layer, categorical alignment increases and peaks at approximately 1.4–1.5 s across streams. This indicates that the initial emergence of category-level structure is not restricted to the task-optimised temporal regime.

Differences become clearer at later processing stages. In the second recurrent layer, the 50–200– 400 ms architecture exhibits weaker and less sharply defined alignment than the task-optimised architecture. This pattern persists after multistream integration and classification, where the longer-window architecture reaches a lower peak classifier correlation than the 20–40–80 ms model.

To compare the resulting representational geometries directly, we examined classifier-layer RDMs at the time of maximal categorical alignment. Figure S13 shows the empirical RDMs for both temporal regimes alongside the ideal categorical model.

Both architectures develop a block-diagonal organisation, indicating lower within-category dissimilarity and higher between-category dissimilarity. However, the task-optimised 20–40–80 ms model exhibits a sharper and more coherent block structure than the 50–200–400 ms comparison architecture. This result is consistent with the RSA time courses and indicates that temporal-integration scales affect the quality and stability of category-level geometry.

One possible explanation is that longer temporal windows introduce temporal smoothing, thereby reducing the preservation of fine-grained discriminative information in environmental sound recognition. This interpretation remains to be tested explicitly using controlled acoustic stimuli.

**Figure S1:**
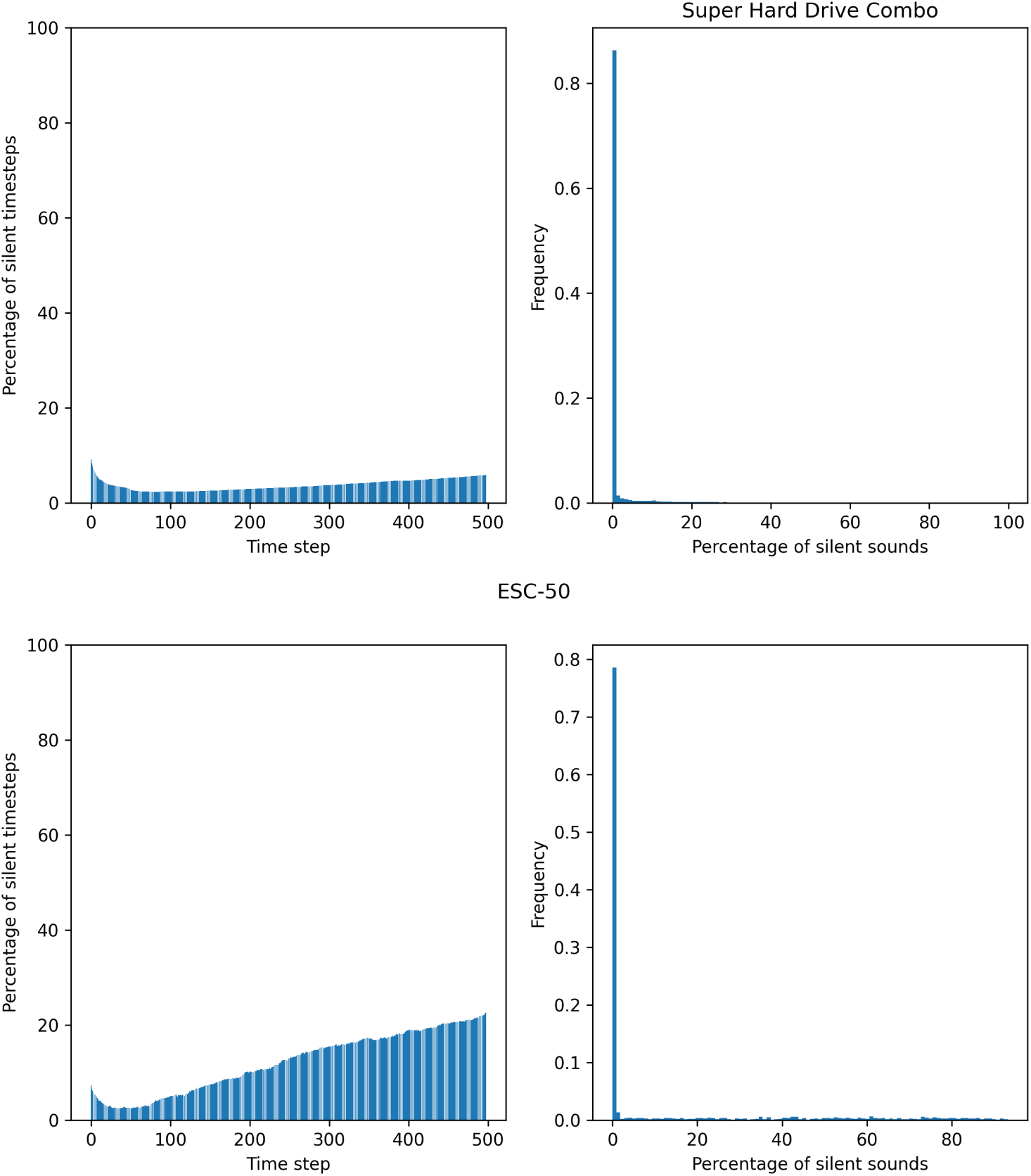
Distribution of silent frames in SHDC and ESC-50. Proportion of silent frames across time and distribution of the percentage of silence per sound. Frames were classified as silent when RMS energy fell below the predefined threshold of 0.1.

**Figure S2:**
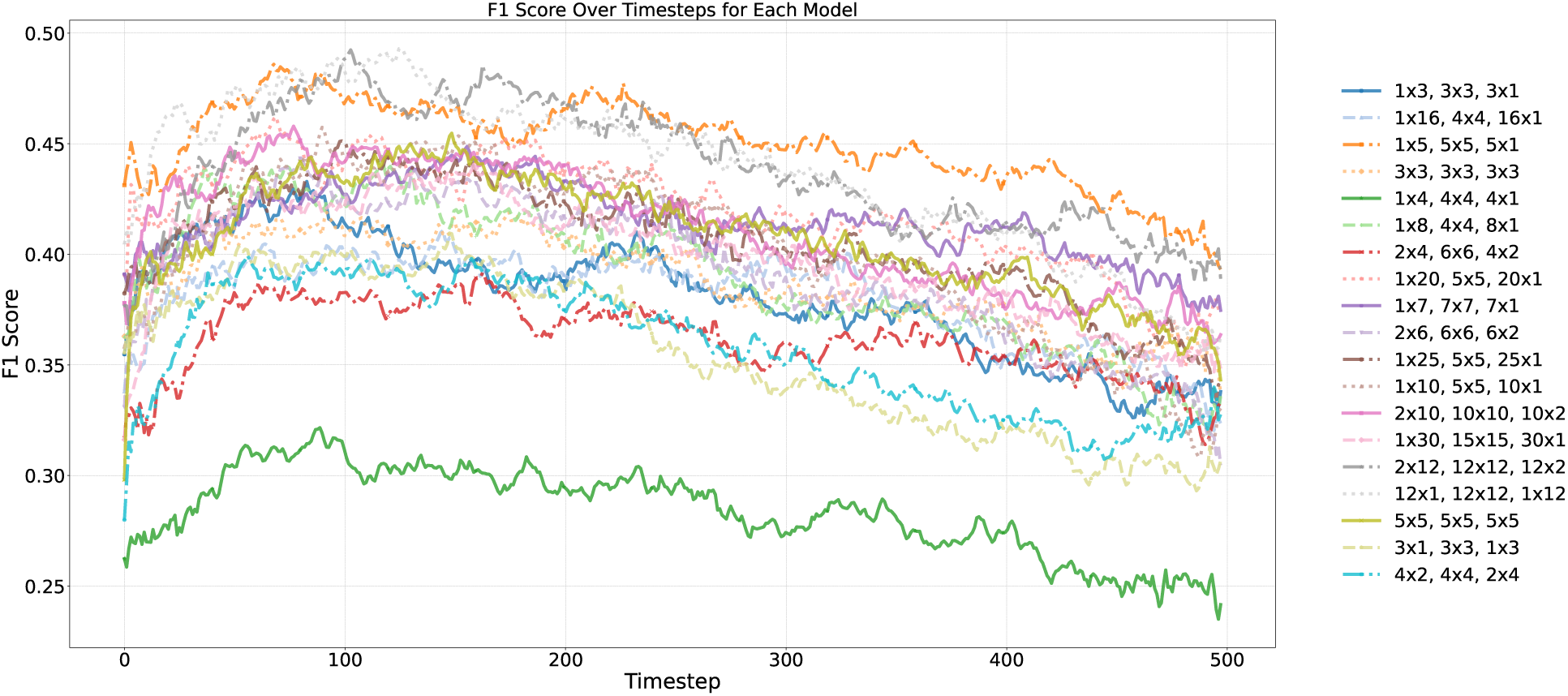
Convolutional-kernel geometry search. Framewise F1 score for alternative convolutional-kernel configurations. The selected configuration used kernels of 1 *×* 5, 5 *×* 5, and 5 *×* 1 for the short, intermediate, and long temporal streams, respectively.

**Figure S3:**
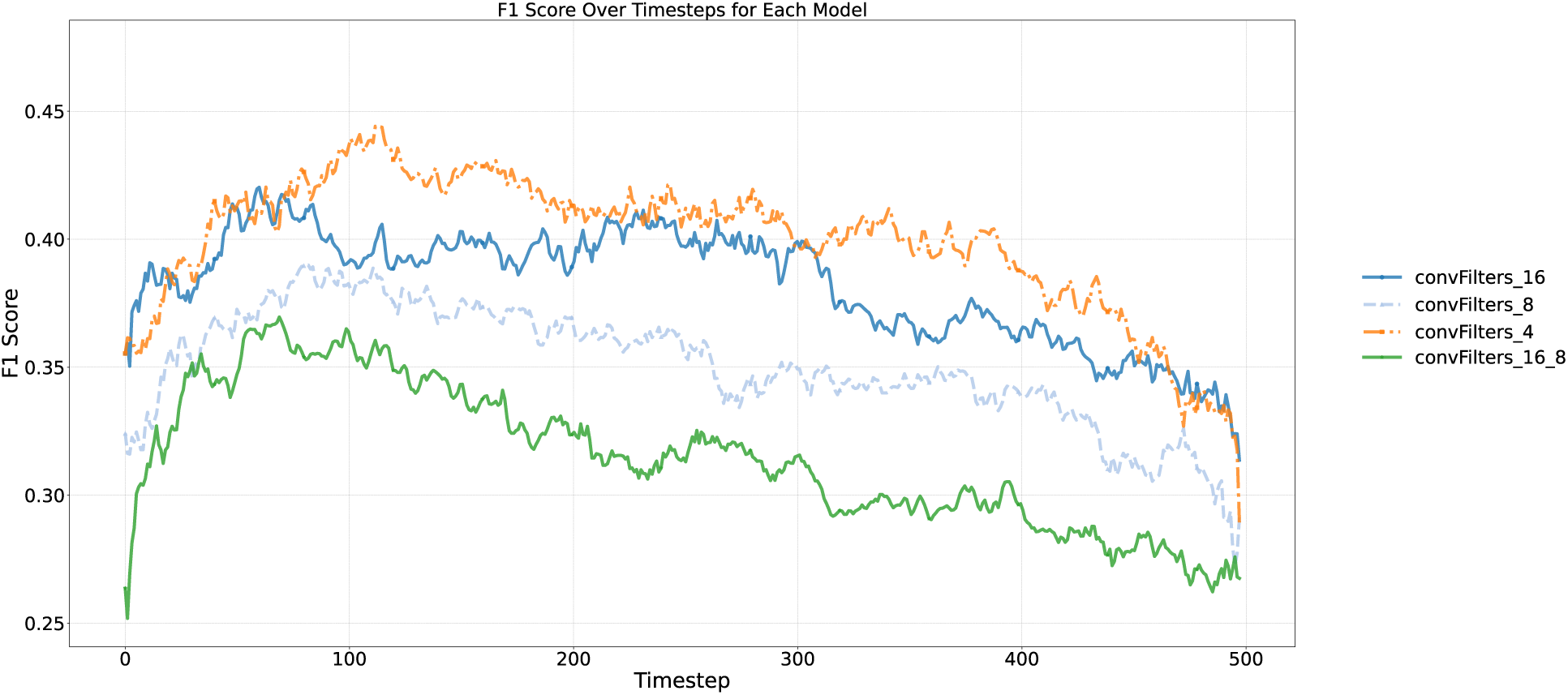
Convolutional-depth and filter-count search. Framewise F1 score for alternative convolutional depths and filter counts. The selected architecture used a single convolutional layer with four filters.

**Figure S4:**
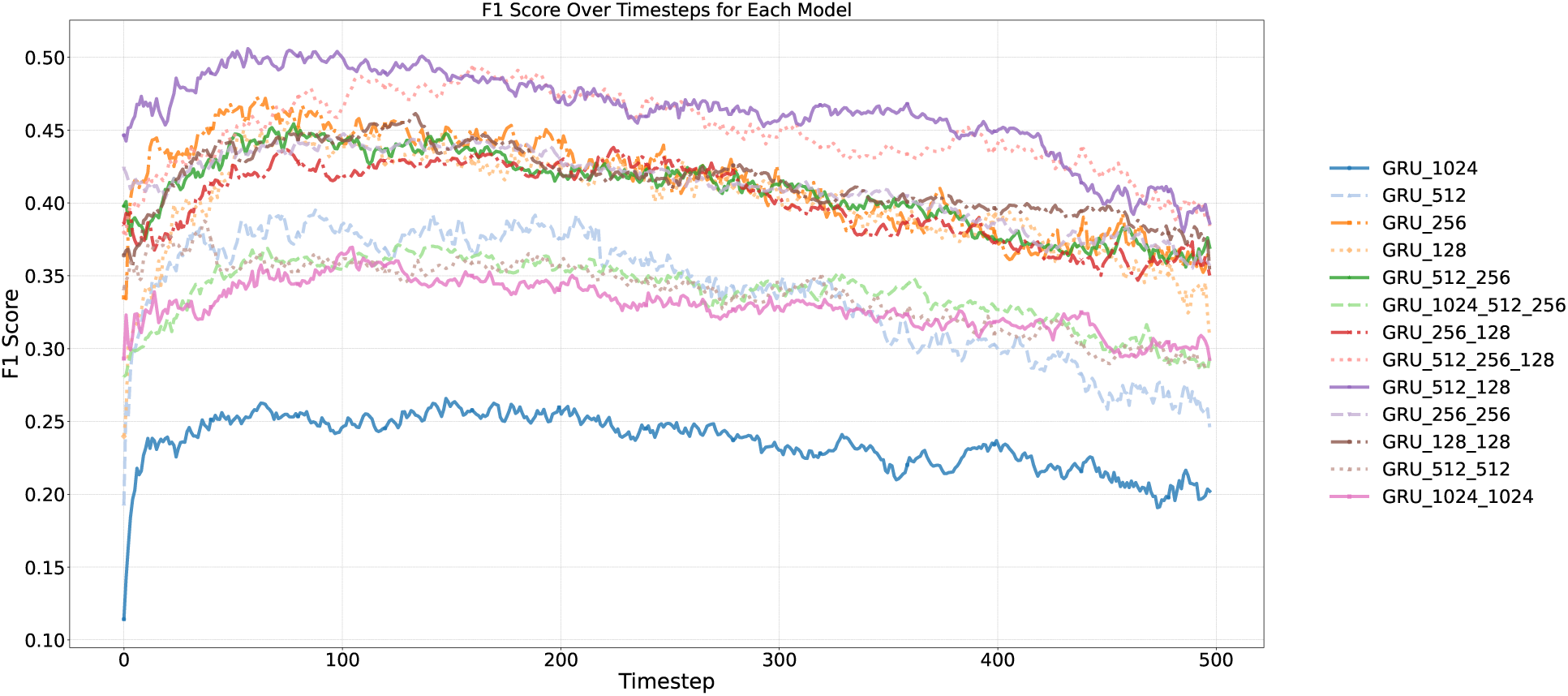
Recurrent-architecture search. Framewise F1 score for alternative GRU depths and dimensionalities. The selected architecture used two GRU layers with 512 and 128 units.

**Figure S5:**
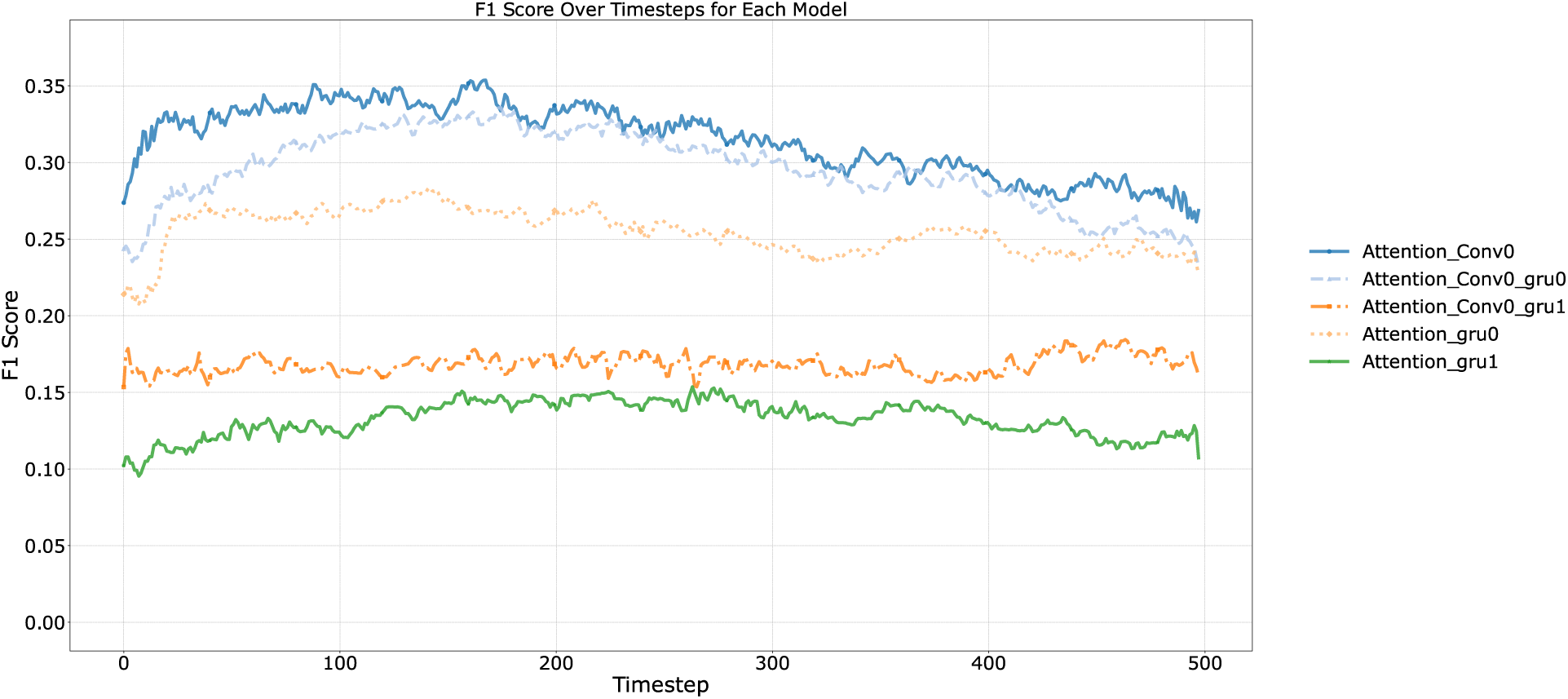
Attention-placement search. Framewise F1 score for alternative attention placements. The selected architecture applied attention after the first convolutional layer.

**Figure S6:**
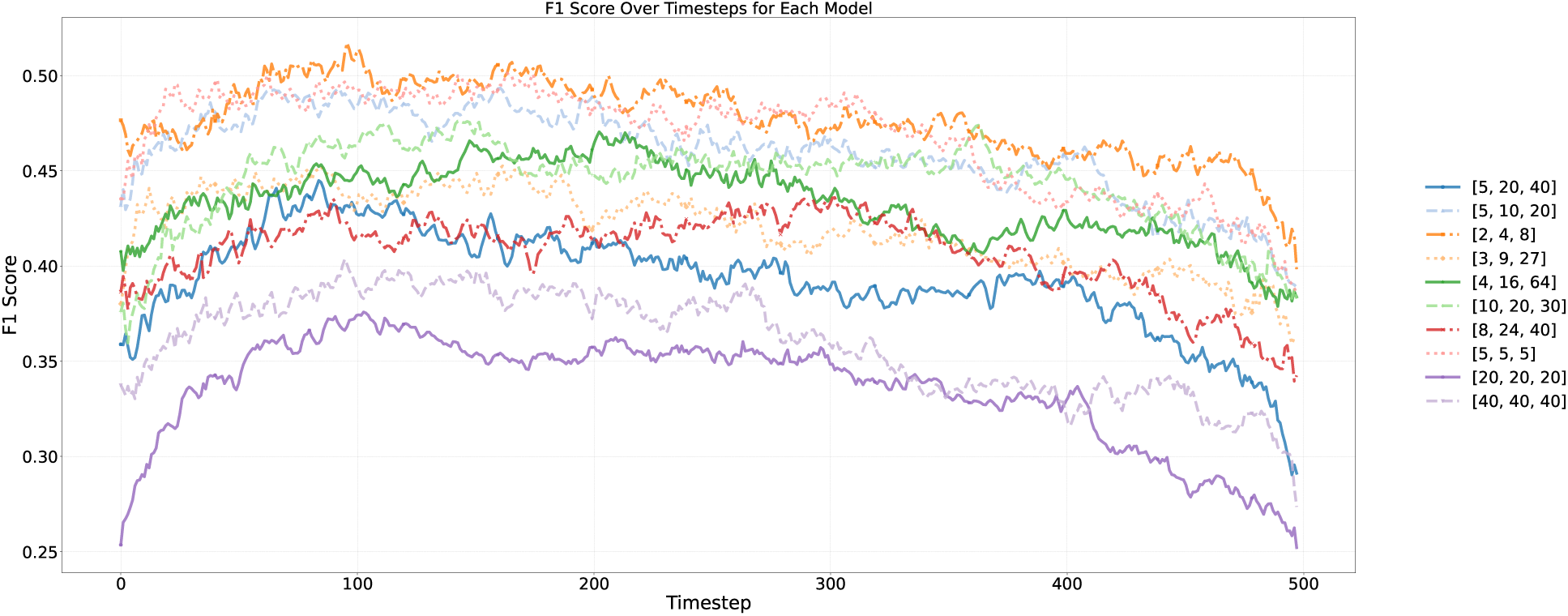
Temporal-buffer configuration search. Framewise F1 score for alternative multiscale temporal-buffer configurations evaluated during architecture calibration. Buffer sizes are expressed in units of 10 ms frames. The selected configuration [2, 4, 8] corresponds to temporal windows of 20, 40, and 80 ms.

**Figure S7:**
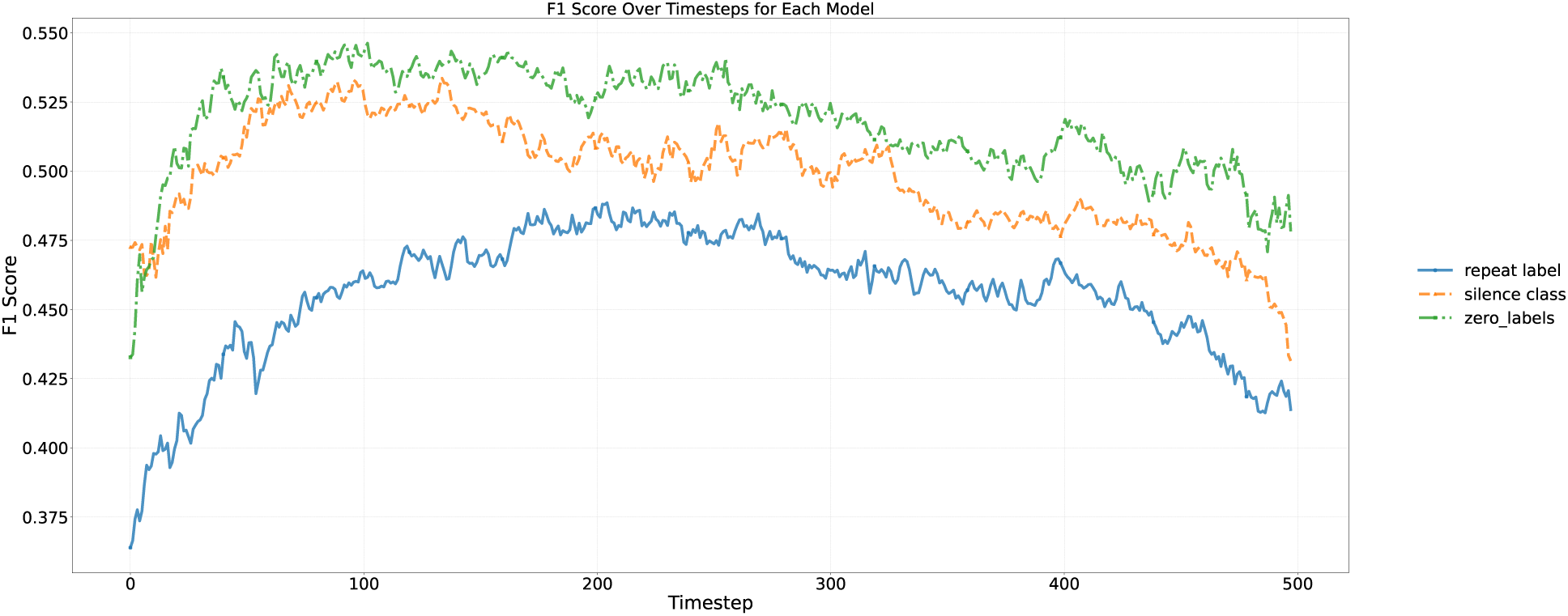
Comparison of silence-handling strategies. Framewise F1 score for models trained using label repetition, a dedicated silence class, or the zero-label strategy for low-energy frames. The zero-label strategy yielded the highest validation performance and was used in subsequent analyses.

**Figure S8:**
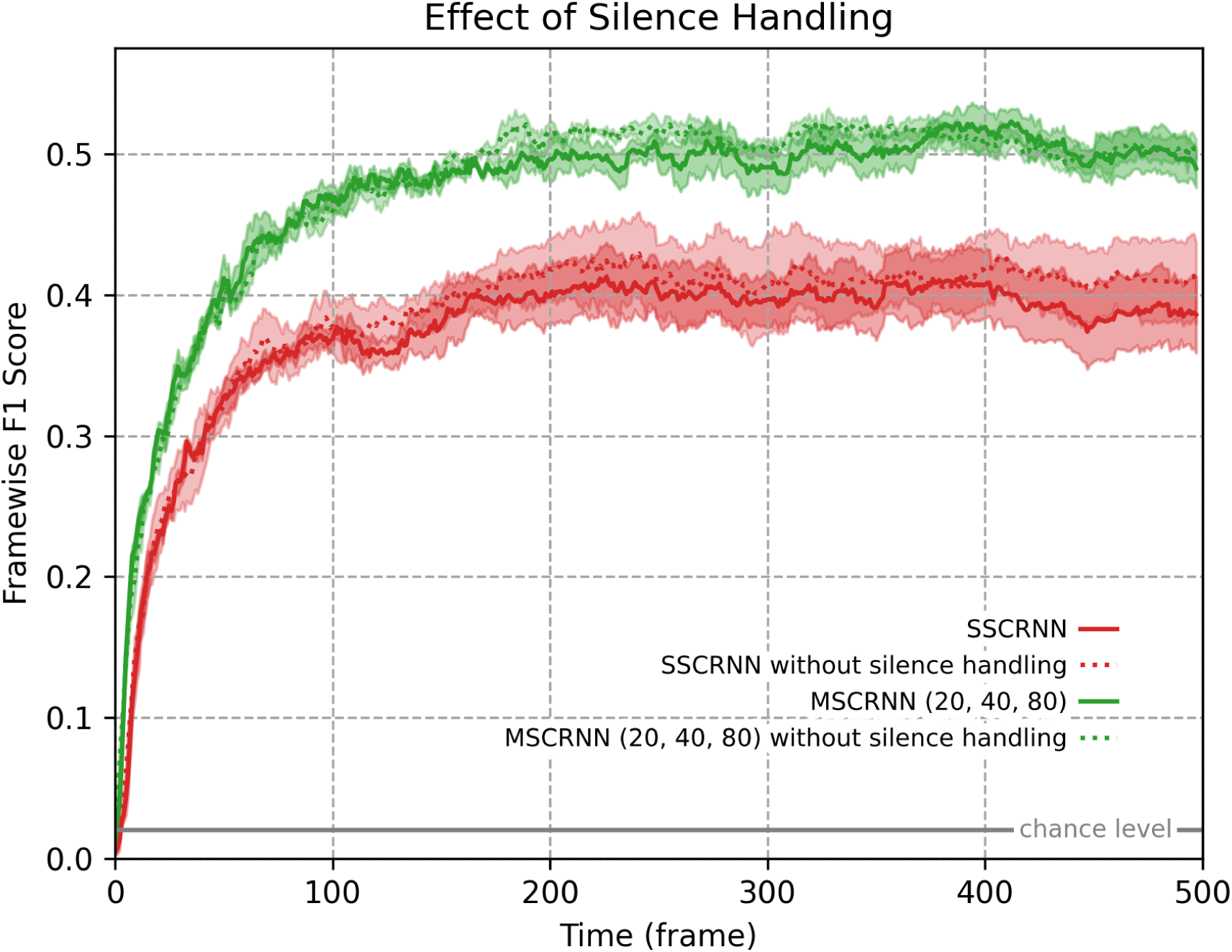
Robustness of model comparisons to silence handling. Framewise F1 score for causal and non-causal single-stream and multistream architectures evaluated without the selected zero-label silence strategy. The qualitative interaction between causality and multiscale processing remains visible.

**Figure S9:**
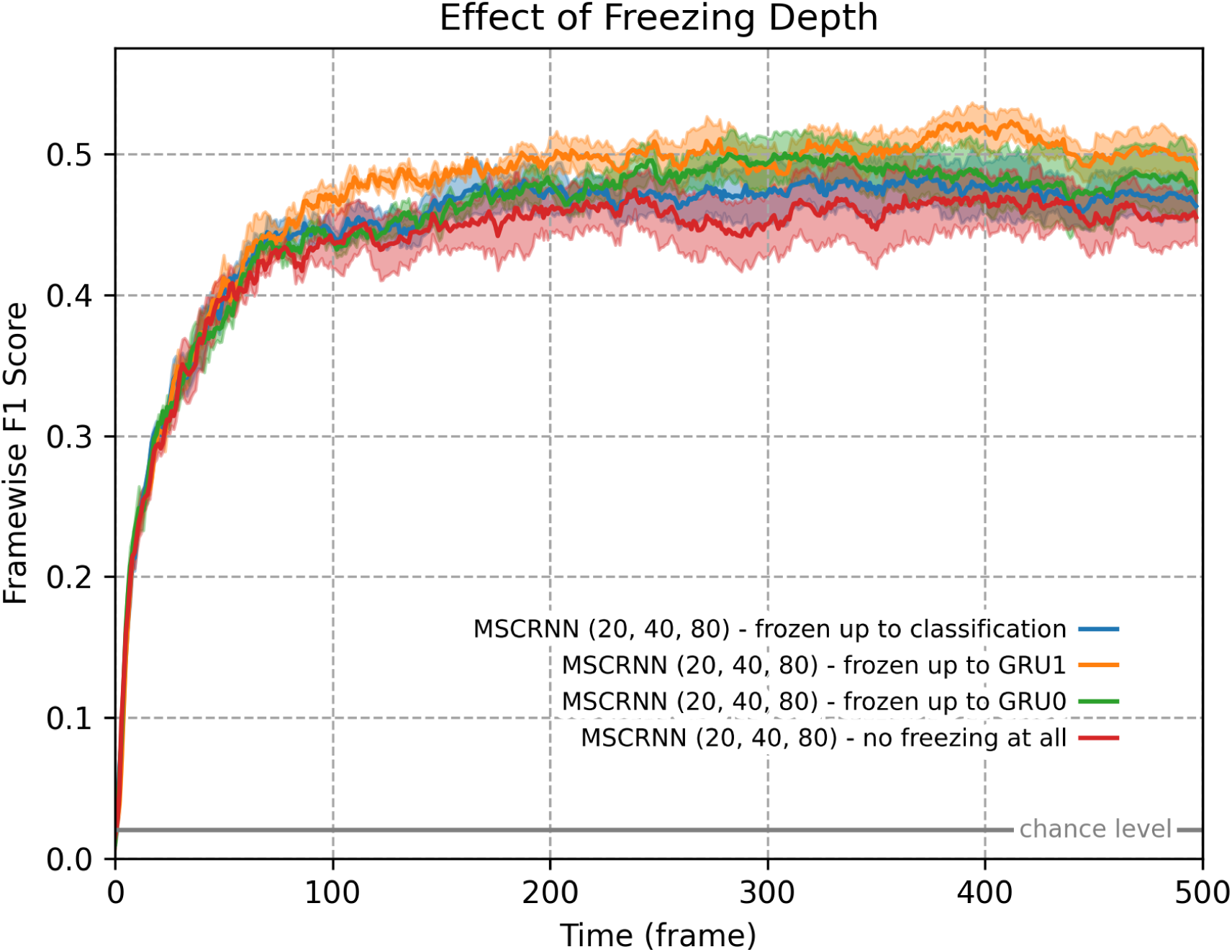
Effect of freezing depth during ESC-50 fine-tuning. Framewise F1 score over time for alternative freezing strategies applied to the task-optimised MsCRNN (20–40–80 ms). Performance was highest when weights were frozen up to GRU1. Shaded regions indicate variability across training runs.

**Figure S10:**
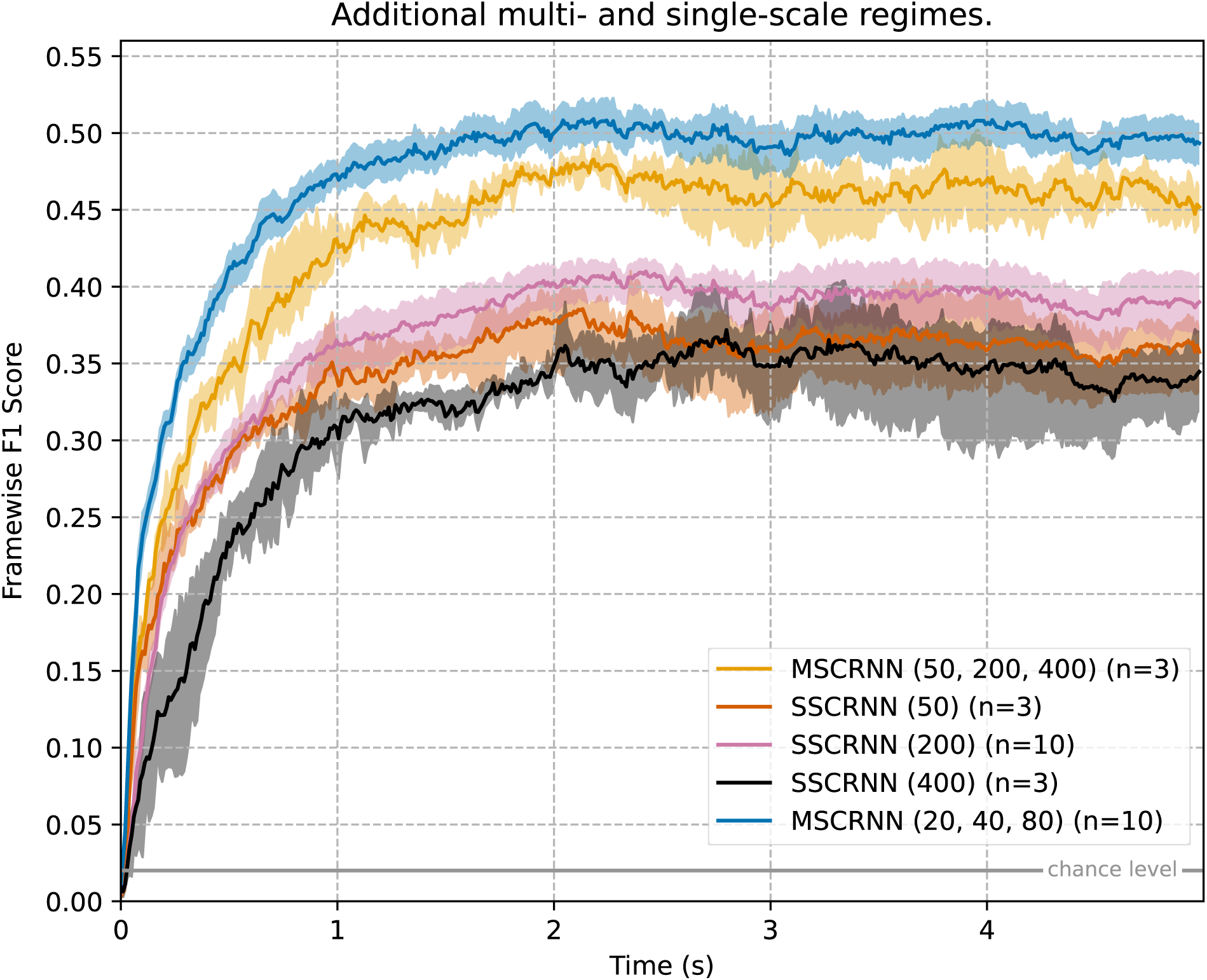
Framewise F1-scores of a 50–200–400ms multistream (MS) and its constituent single-stream (SS) architectures, benchmarked under causal constraints. The multistream configuration consistently outperforms the single stream components. All results report the means of 10 trainings on independent seeds per setting and shaded areas indicate the 95% bias-corrected and accelerated [Efron, 1987] bootstrap confidence intervals (*B* = 10, 000).

**Figure S11:**
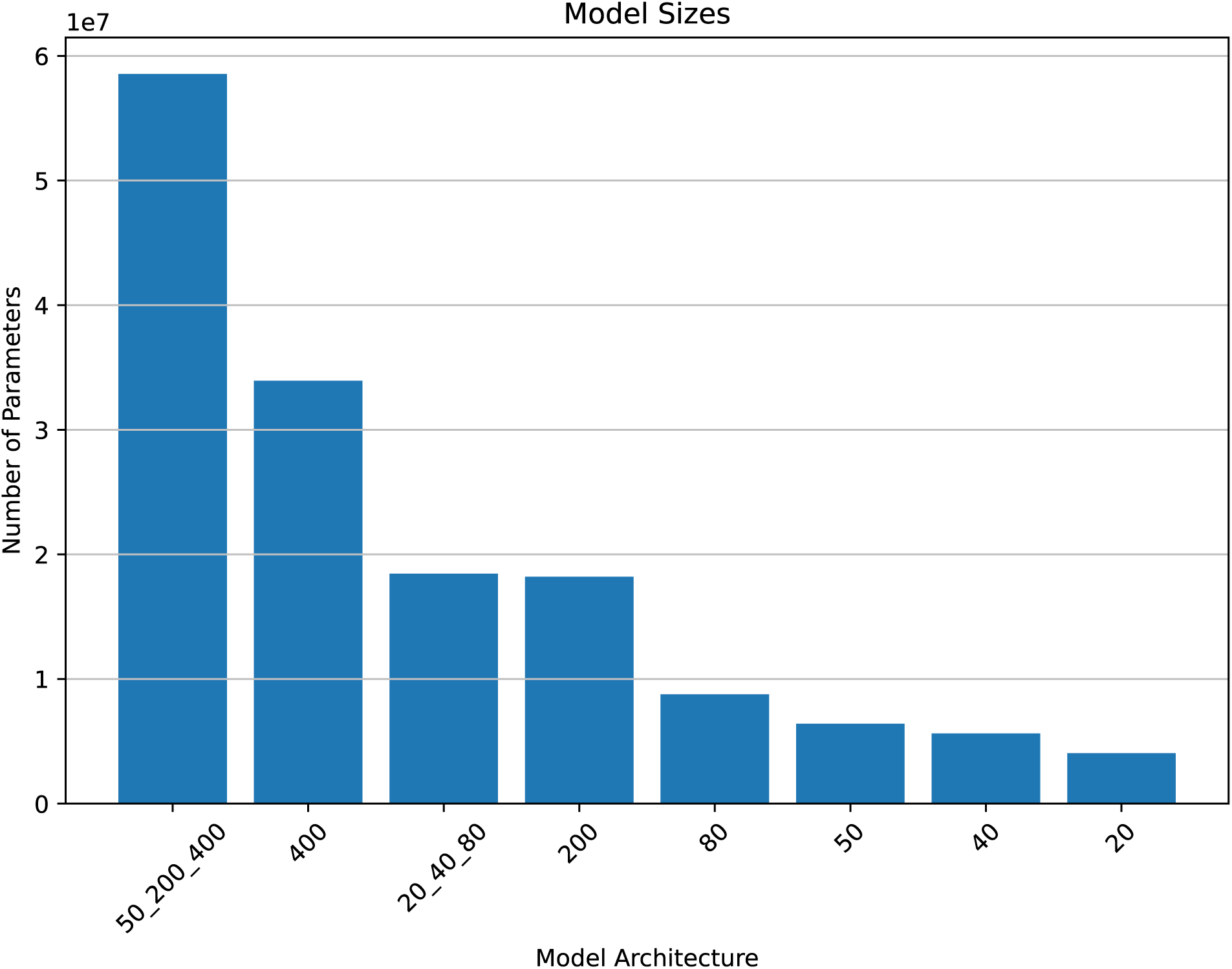
Parameter counts (causal setting) of the two multiscale regimes and all the single-stream constituents.

**Figure S12:**
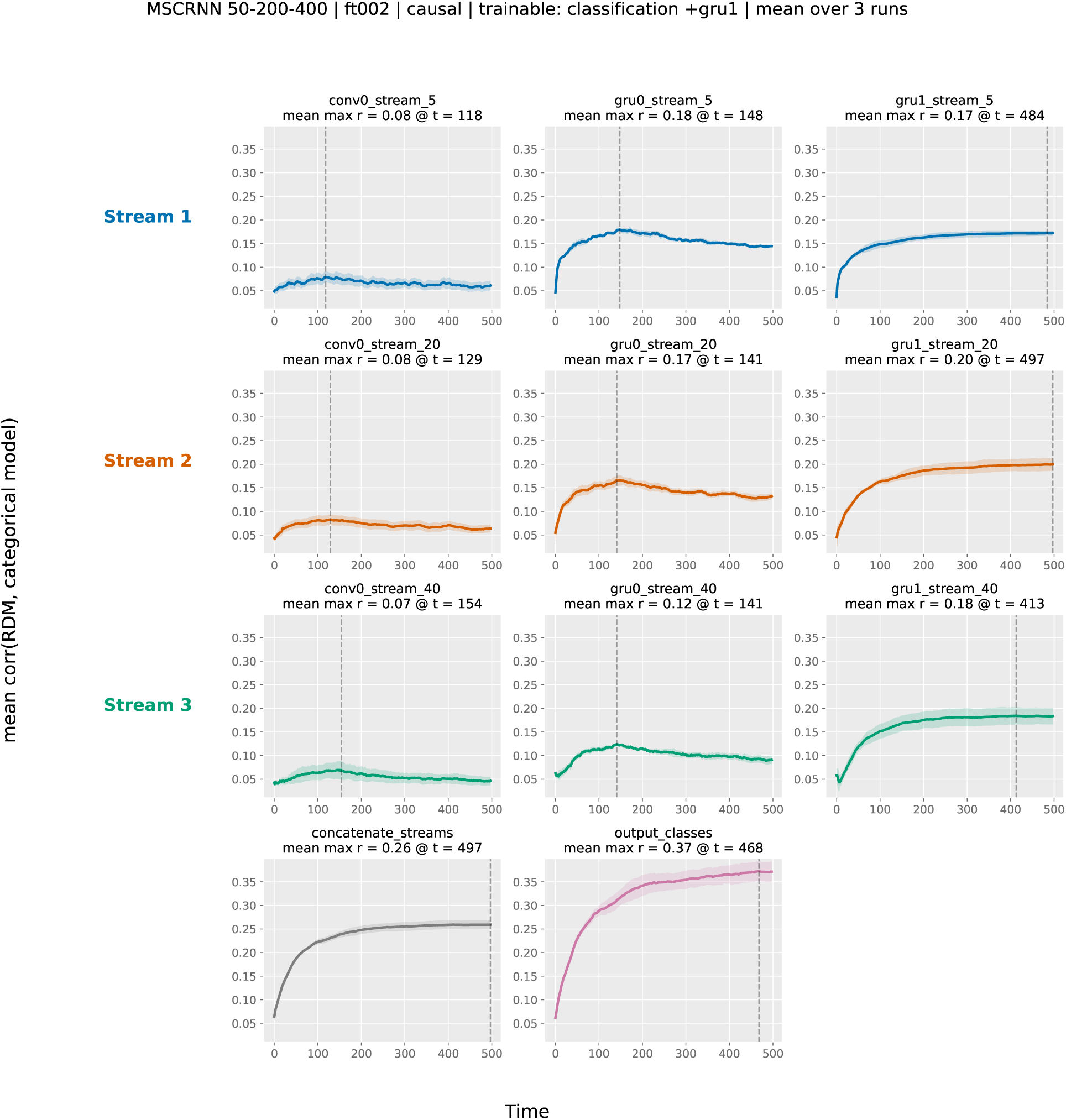
Category-level RSA time courses for the 50–200–400 ms comparison architecture. Time-resolved Pearson correlation between layer-wise RDMs and the categorical model, averaged across training runs. Rows correspond to temporal streams and columns correspond to processing stages. The longer-window architecture develops category-level structure across recurrent processing but reaches lower classifier-level alignment than the task-optimised 20–40–80 ms architecture.

**Figure S13:**
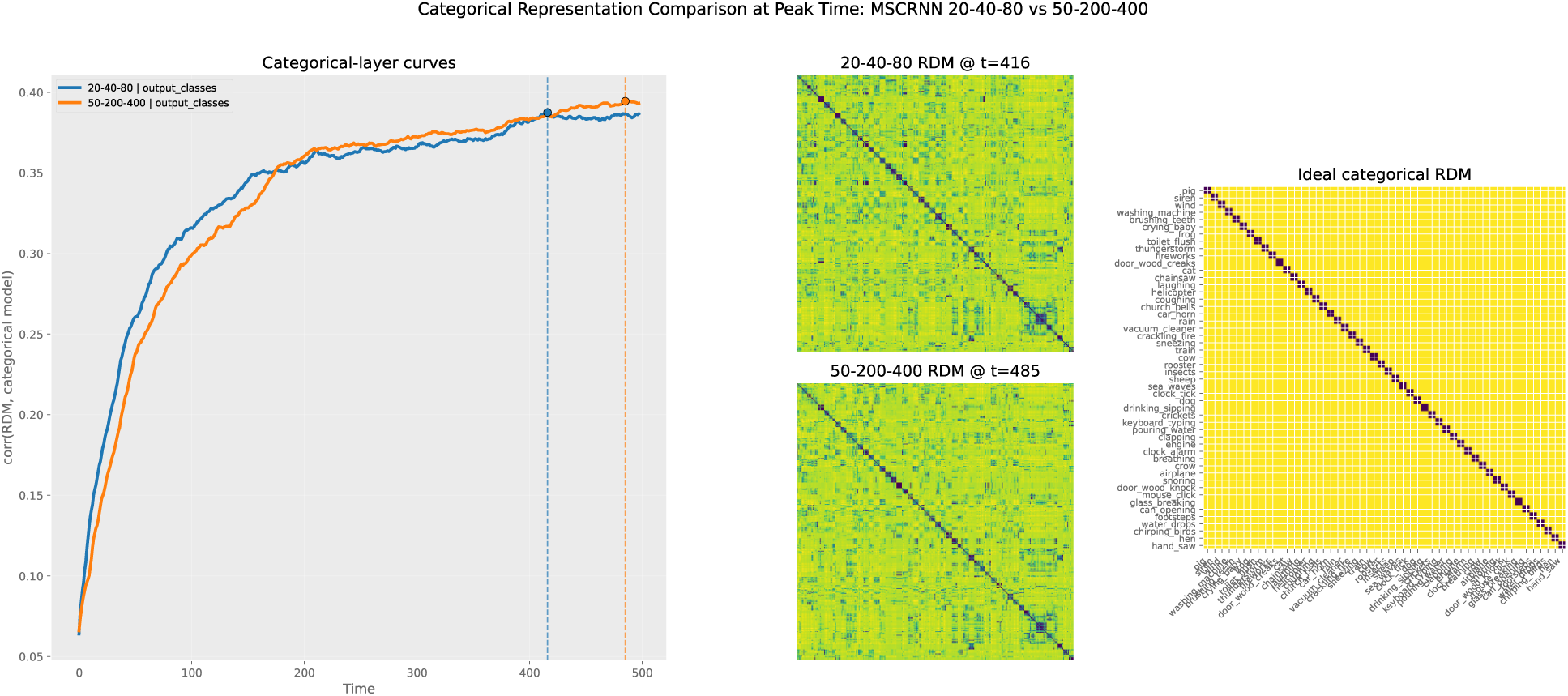
Classifier-level categorical geometry across temporal-integration regimes. Time-resolved categorical alignment and empirical classifier-layer RDMs at peak alignment for the task-optimised 20–40–80 ms model and the 50–200–400 ms comparison architecture. The ideal categorical RDM assigns dissimilarity 0 to pairs from the same class and dissimilarity 1 to pairs from different classes. Both architectures develop category-level organisation, but the task-optimised model exhibits a sharper block-diagonal geometry.

## 3 Supplementary Tables

**Table S2:**
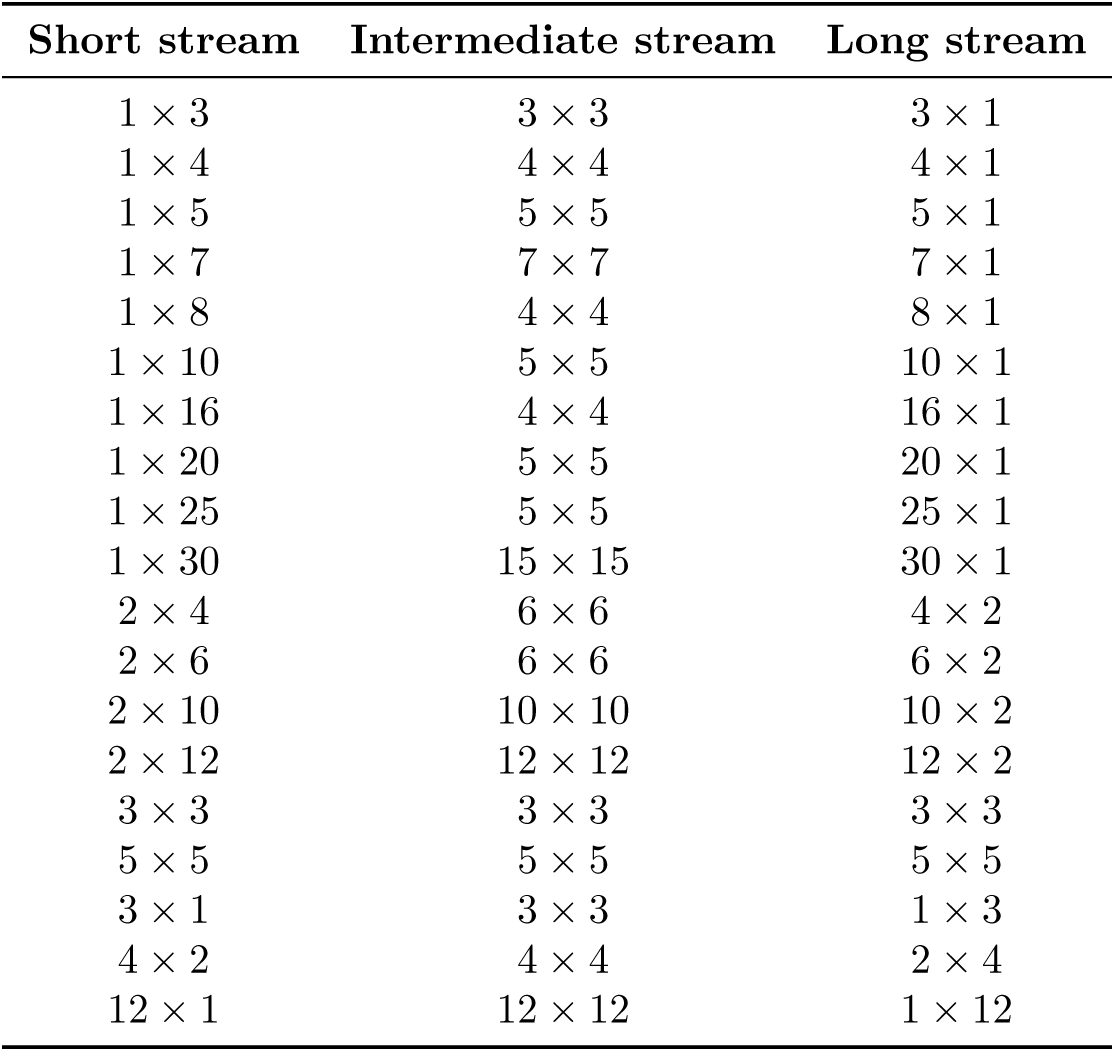
Convolutional-filter geometries evaluated during architecture calibration. Kernel dimensions are reported as time *×* frequency. Control configurations include square kernels and reversed time–frequency layouts.

| Short stream | Intermediate stream | Long stream |
| --- | --- | --- |
| $1 \times 3$ | $3 \times 3$ | $3 \times 1$ |
| $1 \times 4$ | $4 \times 4$ | $4 \times 1$ |
| $1 \times 5$ | $5 \times 5$ | $5 \times 1$ |
| $1 \times 7$ | $7 \times 7$ | $7 \times 1$ |
| $1 \times 8$ | $4 \times 4$ | $8 \times 1$ |
| $1 \times 10$ | $5 \times 5$ | $10 \times 1$ |
| $1 \times 16$ | $4 \times 4$ | $16 \times 1$ |
| $1 \times 20$ | $5 \times 5$ | $20 \times 1$ |
| $1 \times 25$ | $5 \times 5$ | $25 \times 1$ |
| $1 \times 30$ | $15 \times 15$ | $30 \times 1$ |
| $2 \times 4$ | $6 \times 6$ | $4 \times 2$ |
| $2 \times 6$ | $6 \times 6$ | $6 \times 2$ |
| $2 \times 10$ | $10 \times 10$ | $10 \times 2$ |
| $2 \times 12$ | $12 \times 12$ | $12 \times 2$ |
| $3 \times 3$ | $3 \times 3$ | $3 \times 3$ |
| $5 \times 5$ | $5 \times 5$ | $5 \times 5$ |
| $3 \times 1$ | $3 \times 3$ | $1 \times 3$ |
| $4 \times 2$ | $4 \times 4$ | $2 \times 4$ |
| $12 \times 1$ | $12 \times 12$ | $1 \times 12$ |

**Table S3:** Convolutional-depth and filter-count configurations evaluated during architecture calibration. Kernel geometries were fixed according to the convolutional-kernel search.

| Stream | Depth | Filter counts |
| --- | --- | --- |
| Short stream | 1 layer | [4], [8], [16] |
| Short stream | 2 layers | [16, 8] |
| Intermediate stream | 1 layer | [4], [8], [16] |
| Intermediate stream | 2 layers | [16, 8] |
| Long stream | 1 layer | [4], [8], [16] |
| Long stream | 2 layers | [16, 8] |

**Table S4:** GRU architectures evaluated during architecture calibration. Single-layer, heterogeneous multi-layer, and homogeneous multi-layer configurations were compared.

| GRU configurations |
| --- |
| Single layer: [1024], [512], [256], [128] |
| Two layers: [512, 256], [512, 128], [256, 128] |
| Three layers: [1024, 512, 256], [512, 256, 128] |
| Homogeneous two layers: [128, 128], [256, 256], [512, 512], [1024, 1024] |

**Table S5:** Attention-placement configurations evaluated during architecture calibration. Attention placement was kept consistent across temporal streams.

| Attention placement |
| --- |
| No attention |
| After first convolutional layer (conv0) |
| After first recurrent layer (GRU0) |
| After second recurrent layer (GRU1) |
| Spectral attention after conv0 and temporal attention after GRU0 |
| Spectral attention after conv0 and temporal attention after GRU1 |

**Table S6:** Temporal-buffer configurations evaluated during architecture calibration. Buffer sizes are reported in milliseconds and include shorter-scale, logarithmic, neuroscientifically motivated, and equal-window control configurations.

| Temporal-buffer sizes (ms) |
| --- |
| 50 – 200 – 400 |
| 50 – 100 – 200 |
| 20 – 40 – 80 |
| 30 – 90 – 270 |
| 40 – 160 – 640 |
| 80 – 240 – 400 |
| 100 – 200 – 300 |
| 50 – 50 – 50 |
| 200 – 200 – 200 |
| 400 – 400 – 400 |

**Table S7:** Silence-handling strategies evaluated during architecture calibration. Strategies differ in how silent or low-energy frames contribute to training.

| Strategy | Description |
| --- | --- |
| Label repetition | The global sound label is assigned to all frames, including silent frames. |
| Dedicated silence class | Silent frames are assigned to an additional silence category. |
| Zero-label strategy | The target vector is set to zero for silent frames. These frames contribute no class-specific cross-entropy term while remaining included in the loss reduction across time steps. |

